# Engineering a virus-like particle to display peptide insertions using an apparent fitness landscape

**DOI:** 10.1101/2020.06.23.168070

**Authors:** Stephanie A. Robinson, Emily C. Hartman, Bon C. Ikwuagwu, Matthew B. Francis, Danielle Tullman-Ercek

**Affiliations:** Department of Chemistry, University of California, Berkeley, California 94720-1460; Department of Chemical and Biological Engineering, Northwestern University, 2145 Sheridan Road, Technological Institute E136, Evanston, Illinois 60208-3120; Materials Sciences Division, Lawrence Berkeley National Laboratories, Berkeley, California 94720-1460

**Keywords:** self-assembling proteins, peptide insertions, apparent fitness landscape, epitope display, virus-like particle, MS2

## Abstract

Peptide insertions in the primary sequence of proteins expand functionality by introducing new binding sequences, chemical handles, or membrane disrupting motifs. With these properties, proteins can be engineered as scaffolds for vaccines or targeted drug delivery vehicles. Virus-like particles (VLPs) are promising platforms for these applications since they are genetically simple, mimic viral structure for cell uptake, and can deliver multiple copies of a therapeutic agent to a given cell. Peptide insertions in the coat protein of VLPs can increase VLP uptake in cells by increasing cell binding, but it is difficult to predict how an insertion affects monomer folding and higher order assembly. To this end, we have engineered the MS2 VLP using a high-throughput technique, called Systematic Mutagenesis and Assembled Particle Selection (SyMAPS). In this work, we applied SyMAPS to investigate a highly mutable loop in the MS2 coat protein to display 9,261 non-native tripeptide insertions. This library generates a discrete map of three amino acid insertions permitted at this location, validates the FG loop as a valuable position for peptide insertion, and illuminates how properties such as charge, flexibility, and hydrogen bonding can interact to preserve or disrupt capsid assembly. Taken together, the results highlight the potential to engineer VLPs in systematic manner, paving the way to exploring the applications of peptide insertions in biomedically relevant settings.

## INTRODUCTION

The applications for virus-like particles (VLPs) are diverse in the biomedical and nanotechnology fields, and have been generally accepted as new technologies for vaccine and targeted drug delivery research.^1,2^ One of the most well-studied virus-like particles is derived from the MS2 bacteriophage. The MS2 VLP is a chemically robust, genetically simple assembly comprising 180 copies of a single coat protein (CP).^3^ This CP can be easily expressed and harvested in its self-assembled form in bacterial systems, such as *Escherichia coli* (*E. coli*), and can be stored for extended periods of time.^4^ This ease of production and storage has led to much interest in repurposing the MS2 VLP as a drug delivery vehicle or vaccine platform, but both of these applications require the assemblies to be engineered in unique ways.^5^ For example, small molecules can be conjugated to the interior, or the VLP can be reassembled around negatively charged cargo for drug delivery.^6–8^ The cargo can then be targeted to cells using binding groups or epitopes attached chemically.^9^

Specifically, VLP-based vaccines require a method of presenting epitopes, or native viral peptides that generate an immune response.^10,11^ For example, the MS2 VLP has been engineered to have comparable immunogenicity to the commercially available Gardesil-9 vaccine in mice.^12^ Epitopes can be presented on MS2 VLPs through two primary methods. In the first, peptides, glycans, or other epitopes can be conjugated to the VLP via chemical methods after expression. This method is proven, but it lacks control over the number of displayed epitopes per VLP. In the second, peptide epitopes are genetically encoded as fusions or insertions in the CP. This technique is known as phage display, which is a well-established technique for many VLPs and permits control over the number of displayed epitope copies through genetically inserting the epitope in the monomer.^13–15^ However, peptide insertions can drastically change the secondary structure of the CP and the ability for monomers to pack into regular icosahedra. Protein engineering offers several strategies for identifying insertion sites that will maintain properly assembled structures.

One promising method for generating peptide insertions is to incorporate them into a loop occurring early in the MS2 CP sequence, but this strategy requires the genetic fusion of monomers to generate stable capsids from dimeric analogs.^16^ These fusion constructs reduce the number of peptides displayed on the surface by half, as capsid assembly only allows one of the monomers to contain the peptide insertion in the fusion dimer. Therefore, it is fruitful to explore other non-fusion alternatives that may increase the diversity of the MS2 VLP peptide insertion sites. We previously developed a protein engineering technique known as Systematic Mutagenesis and Assembled Particle Selection (SyMAPS). SyMAPS mutagenizes the coat protein (CP) of a VLP, selects for assembly, and generates an apparent fitness landscape (AFL). We applied this technique to explore the mutability and principles of higher order assembly for the MS2 VLP.^17–19^

Through this work we identified a region known as the “FG loop” to be especially permissive to mutations. This loop is unique in its conformations and placement in the packing of MS2 monomers. The FG loop extends into the pores of the MS2 VLP and takes on two distinct conformations (Figure 1). The monomers of MS2 pack into symmetric (C/C) and asymmetric (A/B) dimers, which are defined by the secondary structure of the FG loop.^20,21^ Along the quasi-six-fold axis of the VLP, the FG loop is mostly β sheet in character with a hairpin turn, but along the five-fold axis, the FG loop is largely disordered and the hairpin turn is preserved through hydrogen bonding networks with the backbone. This region is flexible and extends into the pore, making it a promising region to examine how a diverse array of peptide insertions affect capsid assembly.

**Figure 1.**
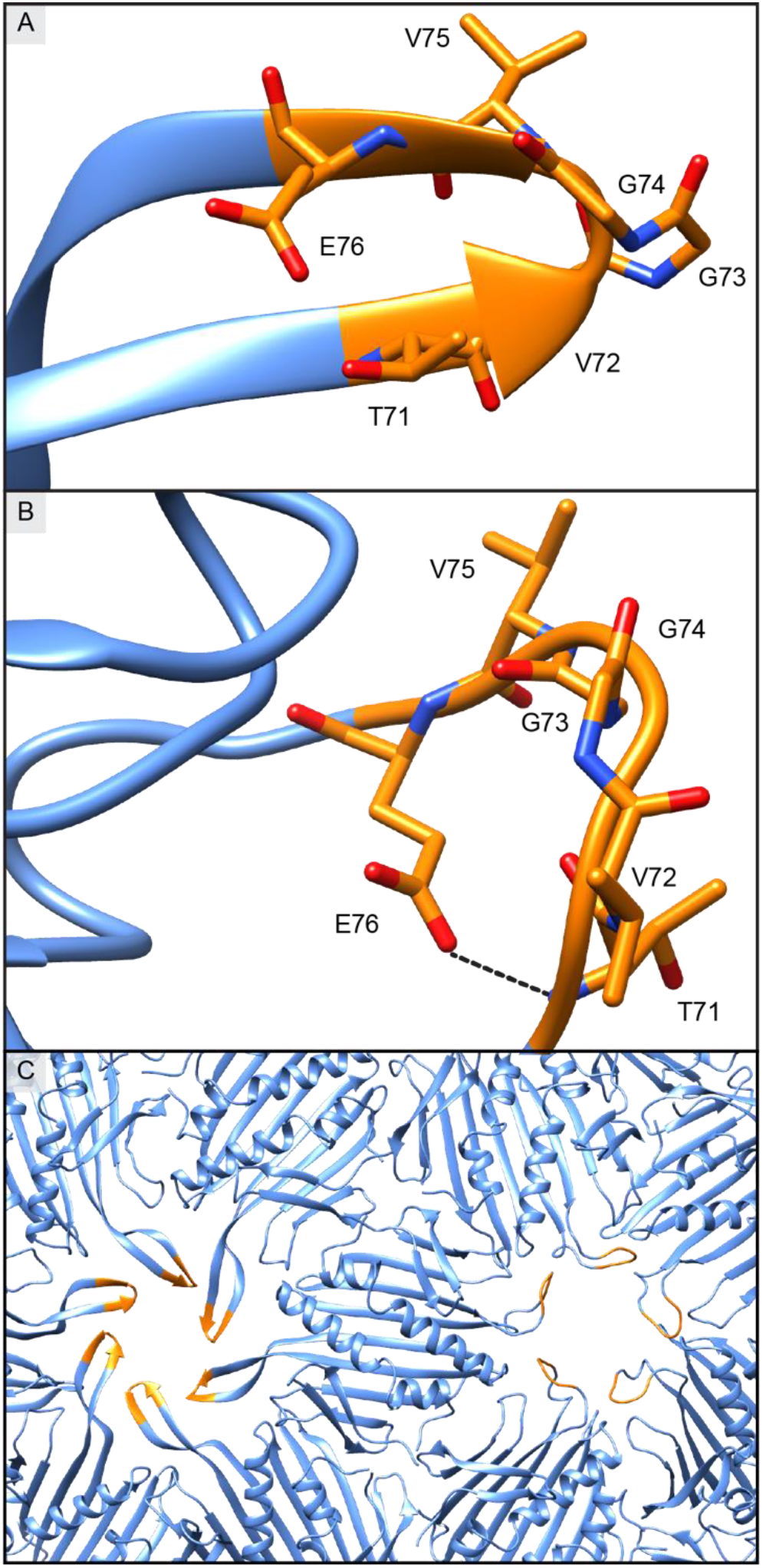
The two conformations of the FG loop. (A) The hairpin turn of the FG loop of a monomer in the quasi-6-fold axis. (B) The disordered FG loop in the five-fold axis. The hydrogen bond between the side chain of E76 and the backbone of T71 is shown by the dashed line. (C) Two conformations of the FG loop assembled into the pores of the MS2 capsid. The quasi-6-fold axis (left) and five-fold axis (right) both highlight the FG loop (residues 71-76) in orange for comparison of the two structures in the monomer packing of the MS2 capsid. PDB 2MS2

**Figure 2.**
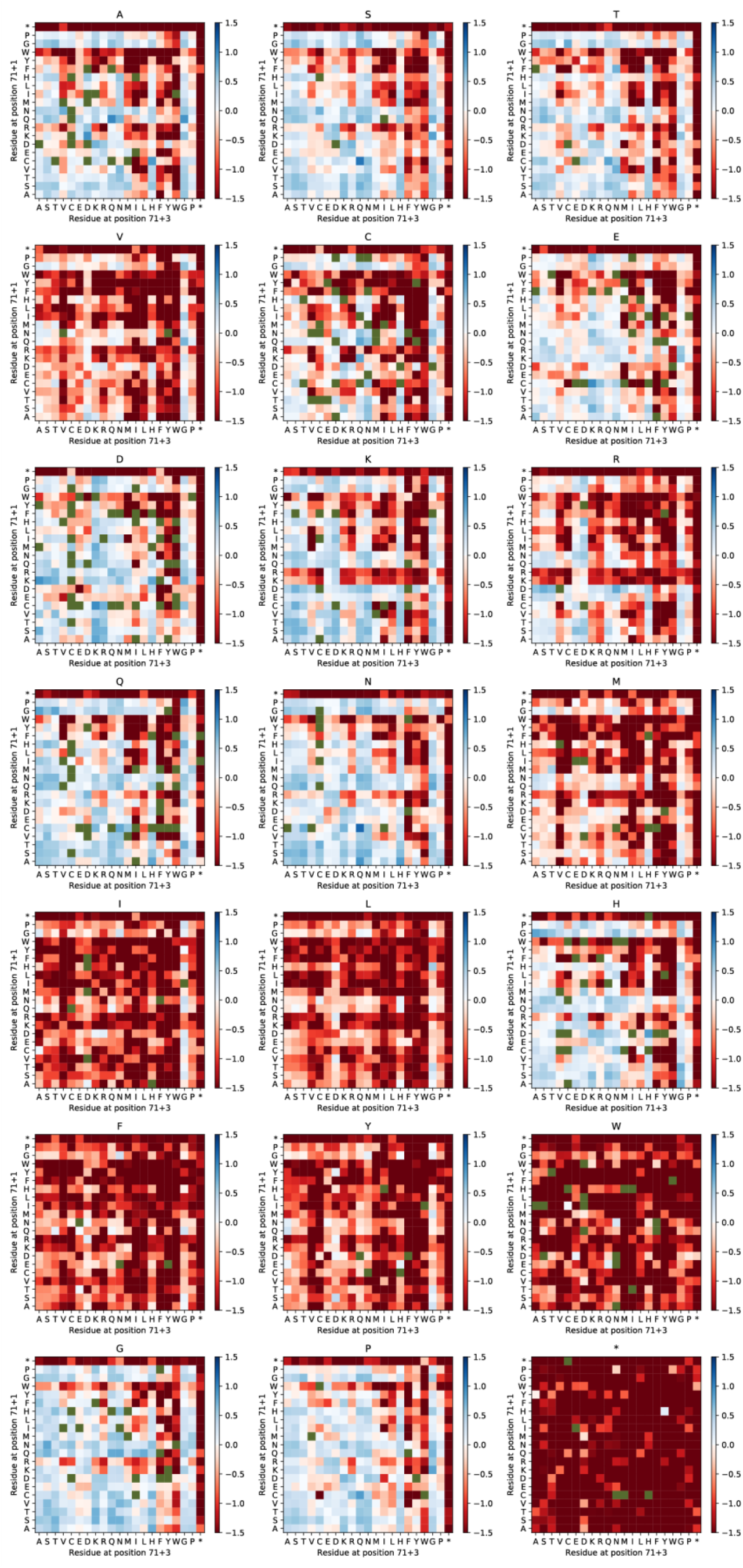
The Apparent Fitness Landscape of the (NNK)_3_ Insertion Library. This AFL is a three-dimensional matrix, so the data are shown as two-dimensional matrices grouped by the identity of the second position. The color of each square corresponds to the AFS assigned to that insertion. A green square is an insertion not sequenced in the plasmid library and is considered a “hole” in the AFL since no score can be assigned. An artificial AFS of −4.0 is assigned to mutants that are present in the plasmid library but not sequenced in the VLP library. Stop codons are labeled with an asterisk (‘*’).

Here, we applied SyMAPS by inserting all possible three-residue peptides in the FG loop of the MS2 CP, selecting for assembly, and sequencing the selected peptide insertion library (Scheme 1). We hypothesized that this loop would be permissive of insertions due to it mutability and flexibility, and the resulting AFL highlights that this region indeed is tolerant of diverse peptide insertions. Moreover, the comprehensive dataset reveals the general properties of specific residues and insertions that are permitted. This study shows the potential for using this variation on our SyMAPS technique to engineer VLPs in a systematic fashion and, specifically, to expand the opportunities for epitope displaying VLPs.

**Scheme 1.**
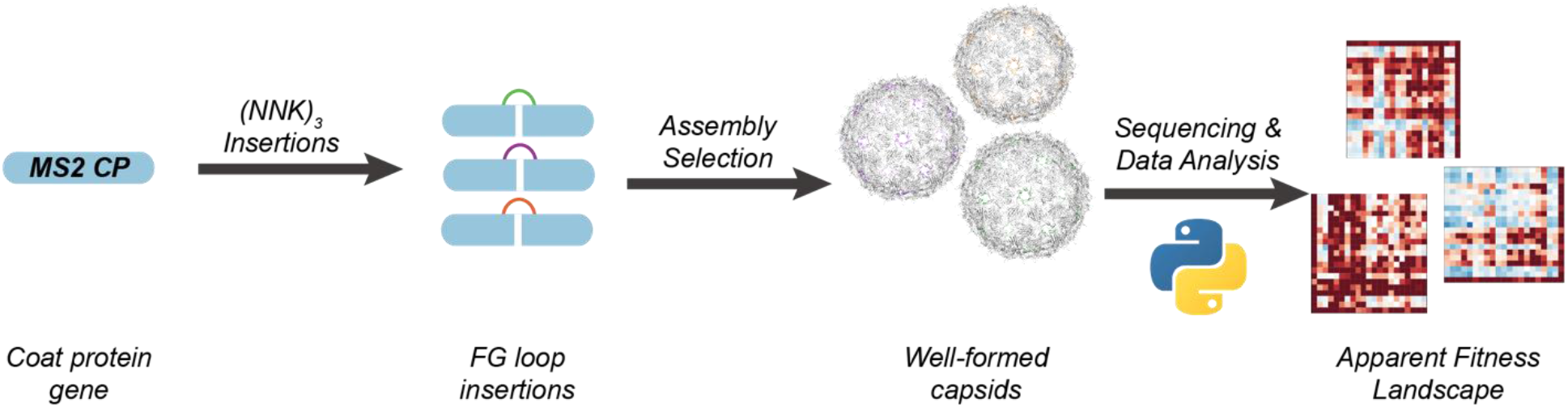
A general scheme for library generation using the SyMAPS method. First, the gene for the MS2 coat protein is diversified with a three-residue insertion library. This library is then expressed, purified, and sequenced to generate an apparent fitness landscape.

## EXPERIMENTAL

### Strains and Plasmids

MegaX DH10B *E. coli* electrocompetent cells (ThermoFisher Scientific, catalog no. C640003) were used for library generation and expression. Chemically competent DH10B *E. coli* were used for individual mutant expressions. The pBAD33 vector was used for all cloning and expression. The key features of this vector are a chloramphenicol resistance cassette and the pBAD promoter, which is induced with arabinose. ^22^

### FG Loop (NNK)_3_ Insertion Library Generation (EMPIRIC CLONING)

The FG Loop (NNK)_3_ insertion library was generated using a modified library preparation strategy developed by the Bolon lab, known as EMPIRIC cloning.^23^ EMPIRIC cloning uses a plasmid with a self-encoded removable fragment (SERF) surrounded by inverted BsaI restriction sites. With this setup, BsaI digestion simultaneously removes both SERF and BsaI sites. These plasmids are termed “entry vectors”, and the SERF in this study encodes constitutively expressed GFP to permit green/white colony screening. This library used a previously generated entry vector that placed a SERF across 26 codons around the FG loop in the MS2 CP.^19^ The fragment that replaces the SERF in the entry vector contains the (NNK)_3_ insertion library. This fragment was synthesized using overlapping forward and reverse single-stranded DNA primers, which were purchased through Sigma-Aldrich. These primers were resuspended in water, pooled, and diluted to final concentrations of 50 ng/μL. The reverse strands were filled in by overlap extension (10 cycles of PCR). The resulting double-stranded DNA fragments were purified using a PCR Clean-up Kit (Promega, Cat# A9282). The purified DNA was then cloned into the entry vector previously described using Golden Gate techniques.^17,23,24^ The ligated plasmids were desalted on membranes (Millipore Sigma, catalog no. VSWP02500) for 20 min. Then the desalted plasmids were transformed into MegaX DH10B *E. coli*, recovered for 1 h at 37 °C with shaking at 200 RPM. The recovered electroporated cells were plated on large (245 × 245 × 20 mm, #7200134, Fisher) LB-agar plates with 32 μg/mL chloramphenicol, and allowed to grow overnight at 37 °C. Then the unscreened library was confirmed to have at least three times the theoretical library size, by plating and counting 1:100 and 1:1000 dilutions of recovered, transformed cells. This process was done in biological triplicate at every step.

### Library Expression and Purification

The three replicates of the colonies for each library were individually scraped from their plates into 10 mL of LB-Miller with 32 μg/mL chloramphenicol and grown for 2 h at 37 °C with shaking at 200 RPM. Each replicate was then subcultured into 1 L of 2×YT (Teknova, catalog no. Y0210) with 32 μg/mL chloramphenicol. The cultures were grown to an OD600 of 0.6, and then expression was induced with 0.1% w/v arabinose. The FG loop (NNK)_3_ insertion libraries were expressed at 37 °C with shaking at 200 RPM overnight (16 h). The cells were then harvested by centrifugation at 5,000 × g for 10 min, resuspended in 10 mM sodium phosphate buffer (pH 7.2) with 2 mM sodium azide, and sonicated at 50% amplitude, pulsing 2 s on and 4 s off for a total of 10 min of “on” time (Fisher Scientific, catalog no. FB120A220, probe CL-18). Do take care when dealing with sodium azide as it is hazardous and explosive when heated. Refer to safety data sheets before handling. Note that from harvesting onwards the libraries were kept at 4 °C. The libraries were precipitated overnight with 50% (w/v) ammonium sulfate and collected by centrifugation at 17,000 × g for 12 min. Pelleted precipitates were then resuspended in sodium phosphate buffer (pH 7.2), 200 mM sodium chloride, and 2 mM sodium azide and syringe filtered (PES 0.45 μm, Fisher Scientific catalog no. 05-713-387).

### FPLC SEC (Assembly Selection)

The FG loop (NNK)_3_ insertion library replicates were purified at 4 °C on an Akta Pure 25 L Fast Protein Liquid Chromatography (FPLC) system with an UV U9-Monitor and a HiPrep Sephacryl S-500 HR column (GE Healthcare Life Sciences, catalog no. 28935607) size exclusion chromatography (SEC) column via isocratic flow with 10 mM sodium phosphate buffer (pH 7.2), 200 mM sodium chloride, and 2 mM sodium azide. Fractions determined to contain MS2 coat protein via sodium dodecyl sulfate–polyacrylamide gel electrophoresis (SDS-PAGE) were collected for further analysis.

### HPLC SEC

The fractions identified to contain well-formed VLP from the assembly selection were combined and run on an Agilent 1290 Infinity HPLC system with a 1290 Diode Array Detector (serial no. DEBAW00292), an Agilent Bio SEC-5 column (5 μm, 2000 Å, 7.8 mm × 300 mm), and an isocratic flow of 10 mM sodium phosphate buffer (pH 7.2), 200 mM sodium chloride, and 2 mM sodium azide.

### High-throughput Sequencing Data Processing

The plasmid library DNA was purified from 2 mL aliquots of cultures before subculturing expressions using a Zyppy Plasmid Miniprep Kit (Zymo, catalog no. D4036). RNA was extracted from the FG loop (NNK)_3_ library following assembly using previously published protocols.^17–19^ In summary, TRIzol (Thermo Fisher, catalog no. 15596026) was used to homogenize collected VLP-containing fractions after assembly selections, and an equal volume of chloroform was added. The samples were separated by centrifugation at 4 °C into aqueous, interphase, and organic layers. The aqueous layer, which contains the RNA from inside the VLPs, was isolated. The RNA was then precipitated with isopropanol and washed with 70% ethanol. The precipitated RNA was then briefly dried and resuspended in RNase free water. cDNA was then synthesized from the isolated RNA using the Superscript III first-strand cDNA synthesis kit from Life (catalog no. 18080051, polyT primer). cDNA and plasmids were both amplified with two rounds of PCR to add barcodes (10 cycles) and the Illumina sequencing handles (8 cycles), respectively, following Illumina 16S Metagenomic Sequencing Library Preparation recommendations. Libraries were combined and analyzed by 150 PE MiSeq in collaboration with the University of California Davis Sequencing Facilities. Reads in excess of 18 million passed filter, and an overall Q30 > 85% was obtained.

### High-Throughput Sequencing Data Analysis

Data were trimmed and processed as previously described with minor variations.^17–19^ Briefly, data were trimmed with Trimmomatic with a four-unit sliding quality window of 20 and a minimum length of 30.

### AFL Calculations

Trimmed high-throughput sequencing reads were analyzed using Python scripts written in-house. Briefly, the mutated region of the MS2 CP was isolated, and the identities of the 3 mutated codons were tallied. Relative percent abundances were calculated as described: the sums of all counts (grand sum) at every combination of amino acids were calculated. We next divided each matrix by its grand sum, generating a matrix of percent abundances. These calculations were repeated for each biological replicate of VLP and plasmid libraries, generating 6 different percent abundance matrices. We then calculated relative percent abundances by dividing the percent abundance for the selected library compared by the percent abundance for the plasmid library for each replicate.

We calculated the mean across the three replicates. All nan (null) values, which indicate variants that were not identified in the plasmid library, were ignored. Scores of zero, which indicate variants that were sequenced in the unselected library but absent in the VLP library, were replaced with an arbitrary score of 0.0001. We calculated the log10 of the relative percent abundance array to calculate the final array for each replicate. Finally, we calculated the average Apparent Fitness Score (AFS) value for each amino acid combination by finding the mean value for every combination.

### Top 1,000 Scoring Mutants Abundances of Residues Calculation

The variants were sorted by decreasing AFS, and the top 1,000 mutants were separated for further analysis. The frequencies of each amino acid and the stop codon in these top 1,000 mutants were counted and the percentages were tabulated. The amino acids were ranked by percent abundance for each position. Then the most abundant residues of each position were recombined and compared with the actual AFS of that variant.

### Individual Insertion Construction, Expression, and Assembly Assay

The 20 insertions identified in Table 1 by concatenating the residues sorted by abundance via the “Top 1000 Scoring Mutants Abundances of Residues Calculation” were individually cloned in a similar fashion to the EMPIRIC Cloning strategy described previously for library generation. Primers for individual insertions can be found in the supplemental information. Individual colonies were then selected and confirmed by submitting for Sanger sequencing. The sequence-confirmed plasmids were then transformed into chemically competent DH10B *E. coli* and recovered in SOC at 37 °C with shaking at 200 RPM. Recovered cells were plated on LB-agar plates with 32 μg/mL chloramphenicol and allowed to grow overnight at 37 °C. Then, 50 mL expressions, harvesting cells, and precipitations were carried out as described previously. The samples were resuspended in 1 mL 1×SEC (pH = 7.2), and syringe filtering with 0.45 μm (Fisher Sci. Cat. 05-713-387) pore size, measuring concentrations via Thermo Scientific NanoDrop 2000c, and normalizing concentrations to 1 mg/mL. Peak heights were measured using size exclusion chromatography using instrument and column. The A280 signal was used to calculate total peak height, subtracting the baseline, and normalizing against the peak height calculated for wild-type (WT) MS2. The assembly assay was then done in triplicate (n = 3) and the values reported are the average peak height with standard deviation as the error bars.

**Table 1.** The abundances of residues in each position of the top 1,000 scoring insertions in the AFL and the concatenated insertions with their corresponding AFS values

| *Position 71+1* |  | *Position 71+2* |  | *Position 71+3* |  | *Concatenated* |  |
| --- | --- | --- | --- | --- | --- | --- | --- |
| Residue | Abundance | Residue | Abundance | Residue | Abundance | Insertion | AFS |
| N | 8.3% | G | 9.6% | G | 10.2% | NGG | 0.470 |
| G | 8.2% | N | 8.8% | N | 8.3% | GNN | 0.750 |
| Q | 8.1% | P | 8.7% | S | 8.0% | QPS | 0.487 |
| S | 8.0% | Q | 8.2% | H | 8.0% | SQH | 0.598 |
| T | 7.1% | S | 7.9% | A | 7.7% | TSA | 0.470 |
| A | 6.8% | K | 7.3% | Q | 7.7% | AKQ | 0.619 |
| P | 6.1% | T | 7.2% | T | 7.6% | PTT | 0.289 |
| C | 5.4% | H | 7.0% | D | 7.1% | CHD | 0.421 |
| D | 5.3% | A | 6.7% | K | 6.5% | DAK | 0.711 |
| H | 5.1% | D | 6.7% | E | 4.9% | HDE | -0.409 |
| K | 5.0% | E | 5.8% | R | 4.6% | KER | 0.182 |
| V | 4.9% | C | 4.5% | C | 4.5% | VCC | -0.712 |
| M | 4.9% | R | 4.5% | P | 3.9% | MRP | -0.686 |
| E | 4.1% | V | 2.2% | M | 3.7% | EVM | -0.515 |
| I | 3.7% | M | 2.1% | V | 3.0% | IMV | -4.000 |
| L | 3.2% | I | 0.8% | I | 1.7% | LII | -0.934 |
| R | 2.8% | L | 0.7% | L | 1.3% | RLL | -1.379 |
| F | 2.0% | Y | 0.7% | Y | 0.8% | FYY | -4.000 |
| Y | 1.1% | F | 0.4% | F | 0.3% | YFF | -4.000 |
| W | 0.1% | W | 0.2% | W | 0.0% | WWW | -1.273 |
| * | 0.0% | * | 0.0% | * | 0.0% | *** | -4.000 |

### Dynamic Light Scattering Measurements

During the HPLC SEC analysis of the Assembly Assay previously described, fractions were collected between 7.3-9.3 min, spin concentrated using MWCO 100 kDa (Fisher Sci Cat. 14-558-404) and diluted to 70 μL at 0.1 mg/mL. Particle sizes measured using Malvern Panalytical Zetasizer Nano ZS (Light Source: He-Ne laser 633nm, Max 5mW) with a protocol specified for protein in water solution. Measurements of a sample were done in triplicate in succession, and the reported values were the calculated Z average diameter and peak 1 average with standard deviation as error bars.

### Thermal Shift Assay

The samples were prepared in a similar fashion to the assembly assay except concentrations were not normalized before collecting fractions (7.3-9.3 min) with a HPLC SEC. Collected fractions were spin concentrated using MWCO 100 kDa (Fisher Sci, catalog no. 14-558-404). Then samples for the thermal shift assay were prepared with final concentrations of 0.5 mg/mL capsid with 20x Sypro Orange Protein Gel Dye (Sigma-Aldrich, catalog no. S5692-50UL). The samples were prepared in triplicate in low profile, white PCR strips (Bio Rad, catalog no. TLS0851) and optical, ultraclear caps (Bio Rad, catalog no. TCS0803). On a Bio Rad C1000 touch Thermal Cycler fitted with a CFX96 Real-Time system the following melting protocol was run: 25 °C, increase temperature 0.5 °C, incubate for 1 min, plate read on “All Channels”, repeat for 110 cycles (or until 80 °C). The Hex channel signal was then exported as a .csv file, and the melting temperature was calculated by finding the minimum of the derivative of the fluorescence signal. Values reported are the average melting temperature (n = 3) with standard deviation as the error.

### Transmission Electron Microscope Imaging

Samples for transmission electron microscopy (TEM) were prepared on carbon/copper grids (Fisher Scientific, catalog no. 50-260-34). Grids were cleaned with a PELCO easiGlow by applying negative charge for 10 sec at 15 mA and 0.26 mBar. Then 10 μL of sample was applied to the grid, allowed to set for 2 min and excess liquid was wicked away with a filter paper. Grids were then washed with 30 μL of Milli-Q water and excess liquid was wicked away. Then, samples were fixed by applying 10 μL of 2% (v/v) glutaraldehyde, allowing to set for 2 min, wicking away the excess liquid, and repeating the washing method with 30 μL of Milli-Q water. Samples were stained with 10 μL of 1% uranyl acetate (w/v), allowed to set for 2 min, and excess liquid was wicked away. Images were taken with a Hitachi HD-2300 Dual EDS Cryo STEM at 200 kV.

### Distribution of AFS Values for Insertions with Multiple Charges

The library was filtered for only insertions that contain at least two of the following residues: D, E, K, and R. The distribution of AFS values were reported as a box and whisker plot and p values were calculated using a pairwise t-test with a two-tailed assumption.

### Calculations for Physical Properties

The values for the physical properties for all amino acids were defined previously in calculations using this same analysis.^17, 25–33^ The physical property values were normalized to between 0 and 1 to allow comparison of relative preference, and the AFS values were normalized between −1 and 1. The tolerance of a given residue for each physical property was obtained by the summation of the apparent fitness score for every amino acid in a position multiplied by its normalized physical property value. Therefore, an overall negative score for residues means the given property is detrimental to fitness and a positive score is well tolerated.

## RESULTS AND DISCUSSION

### Applying SyMAPS to the Systematic FG Loop Peptide Insertion Library

VLPs such as the MS2 bacteriophage offer useful platforms to display epitopes using peptide insertions. We set out to engineer peptide insertions in MS2 in a high through-put manner and assess the tolerance for peptide insertions in the FG loop systematically, and to determine the variety of sequences that the loop might tolerate. To do so, we generated a comprehensive library in a high-throughput manner. Briefly, we generated the tripeptide insertion library between residues 71 and 72 of the MS2 CP—a mutable position early in the FG loop—using the SyMAPS technique.^17^ The insertion library was synthesized using PCR and Golden Gate cloning to insert diversified codons into the MS2 CP gene. We used three diversified NNK codons, so that any nucleotide was allowed for the first two bases but only guanine or thymine was allowed for the final base. This resulted in a codon mixture that encoded for all 20 amino acids while minimizing the number of stops codons in the library. The library was expressed, and well-formed particles were selected through size-exclusion chromatography (SEC), since WT MS2 has a characteristic diameter of 27 nm. Importantly, while the MS2 CP can bind to its genomic RNA, it can similarly encapsulate negatively charged cargo. Well-assembled VLPs will assemble around available negative charge in the cell, including their own mRNA. Since it is unlikely that there will be more than one plasmid or library member per cell, this method establishes a useful genotype-phenotype link, in which nucleic acids are preserved only if the associated variants are well-assembled. Following SEC, the encapsulated RNA from collected fractions was extracted and then sequenced via Illumina high-throughput sequencing. The AFL was then generated by comparing relative abundances of sequences in the plasmid library to the purified VLP library; we call the log10 of this ratio of relative abundances the apparent fitness score (AFS). An enrichment of relative abundance indicates that a certain mutant is well-assembling and results in a high AFS (*i.e.* > 0.2), while de-enrichment or the absence of sequences in the VLP means that mutant is non-assembling or weakly-assembling and results in a low AFS (*i.e.* < 0.2).

### Assessing the Quality of the AFL

We next evaluated the validity and quality of the AFL. We can assess the quality of the data by 1) quantifying the percent saturation of library members in the plasmid library, 2) examining the behavior of the stop codon in the library, and 3) analyzing the reproducibility of apparent fitness scores between the biological replicates.

In order to assess coverage of the 9,261 members (including the stop codons), we examined how many variants were sequenced in the plasmid library. Sufficient saturation of the plasmid library is necessary to have a significant probability that if a variant was not observed in the VLP library, it was expressed but did not assemble into a well-formed capsid. Sequencing the plasmid library revealed 97% of all possible variants in all three replicates. About 3% of the possible mutants were not generated in the plasmid library, and therefore could not appear in the VLP library. Therefore, a significant portion of the library was likely expressed, and the resulting VLP library can be used to infer trends of peptide insertion variants in relation to capsid assembly.

Next, the AFL itself was examined to see if the VLP library exhibited the expected scores when a stop codon was present in the CP variant. Since the insertions were introduced by inserting three codons with random bases, a significant number contain a stop codon. While the (NNK) libraries are designed to minimize the probability that a stop codon will be present in the plasmid library, there will still be some truncated mutants arising from the TAG codons that are included. We used this subset of the library to ensure quality of assembly selections and high-throughput sequencing by examining these mutants as a built-in negative control. Thus, the AFS for each stop codon is included in Figure 3 (labelled with an asterisk in the AFL). The average AFS for insertions containing a stop codon is −3.15. Only 1 of the 1,261 possible variants with a stop codon had an apparent fitness landscape score above zero, and 69% of these variants were screened out of the VLP library entirely (AFS = −4.0).

**Figure 3.**
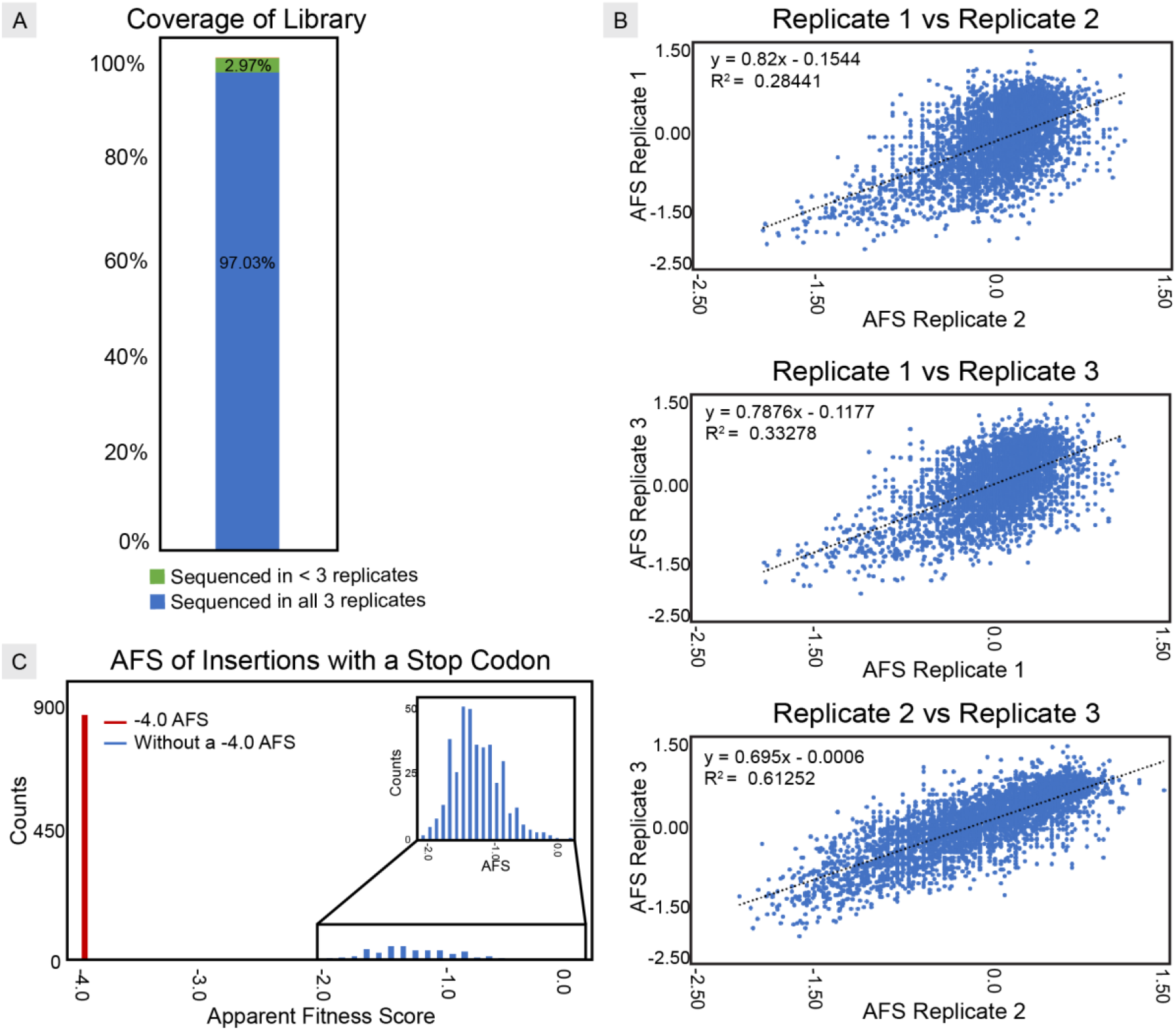
Three methods for checking the quality the of the library data generated. (A) A bar graph showing the coverage of the library in the plasmid library generated over three replicates. We sequenced 97.03% of the library in all three replicates of the plasmid library. Library members not sequenced in one or more replicate of the plasmid library have no AFS and are “holes” in the AFL. (B) The AFS values in each replicate are compared to examine the consistency of the library across replicates. (C) The distribution of AFS values of insertions with a stop codon shows that stop codons were broadly assigned negative AFS values in the AFL.

Lastly, the library should be consistent between each biological replicate. The library was generated in triplicate to minimize noise and random biases in a single sample. To examine how much variance there is between each replicate library, the AFS from each variant in a sample were plotted against the same variant’s score in another replicate and a linear trend was fit to the data. Perfect correlation of these scores would yield a fit with an R^2^ value of 1. We observed R^2^ values from 0.28 to 0.61. These comparisons showed that, while replicate 1 correlated less well with replicates 2 and 3 (R^2^ = 0.28, 0.33 when compared to replicates 2 and 3 respectively), replicates 2 and 3 did correlate well (R^2^ = 0.61). These R^2^ values were consistent with previously published MS2 libraries that used the same SyMAPS technique.^17–19^ Though there is some experimental noise, there is an observed positive correlation between the scores of the variants in one replicate to the respective scores of the variants in another replicate.

The FG loop (NNK)_3_ insertion library quantified here ultimately has the attributes of a high-quality dataset: sufficient saturation of the plasmid library, expected behaviors for the stop codons, and a positive correlation between biological triplicates. Error and noise in these data can be explained by statistical coincidences and compounding random noise, as well as other factors such as variance in expression and cell growth. This noise is subsequently mitigated by quality control of high-throughput sequencing and careful calculation of relative abundances.

### Evaluating General Trends from the AFL

With this dataset in hand, we evaluated overall trends in mutability in the tripeptide insertions. We first grouped the data by the second residue: representing the data in this way can give a visual overview of the trends when the identity of the second position is fixed. We further evaluated the dominant properties that indicate whether a tripeptide insertion is tolerated. We observed that the ability to hydrogen bond and balancing charges were favored in the library. Conversely, the properties that were less tolerated are steric bulk and hydrophobicity, which was expected since the FG loop extends into the pore of the MS2 VLP and is solvent accessible. Since both conformations of the FG loop require a hairpin turn, it is intuitive that the best accommodated residues would be small and flexible.

To analyze the trends for each position further, we calculated how frequently any given amino acid appears in the top 1,000 highest-scoring insertions in the AFL. If every amino acid were equally represented, then all twenty amino acids would have a calculated frequency of 5%. Deviations from this expected frequency reflects a bias for or against that amino acid.

Using this method, we calculated the abundances of each amino acid in the top 1,000 scoring variants (Table 1). We observed that hydrogen bonding amino acids are dominant in the top scoring variants. Upon analysis, 60-70% of the top 1,000 scoring variants in the AFL contain hydrogen bonding residues (S, T, E, D, N, Q, P, K) each position, compared to a theoretical 40% in the unbiased case. Close examination of the native structure in WT MS2 reveals that a key hydrogen bond occurs between the backbone of T71 and E76.^19^ We hypothesize that new hydrogen bonds may help stabilize the tripeptide insertion both in the five-fold and six-fold axes of the MS2 VLP, rearranging the structure to accommodate the additional residues.

Residues that commonly disrupt secondary structure are also disproportionally represented in the top 1,000 scoring insertions. In particular, glycine was the most abundant amino acid in every position (~10% abundant in every position in the top 1,000 scoring variants). The favorability of glycine may be a consequence of the less restricted rotation along the phi and psi angles to access hydrogen bonds with the backbone. Interestingly, when examining the top 1,000 scoring mutants, proline was disfavored in the first and third positions but favored in the second position. This may be due to how proline can “pin” a hairpin turn and stabilize the insertion. Since both proline and glycine disrupt the geometry and secondary structure of the protein, we hypothesize that the backbone may be adopting unusual conformations in order to access favorable hydrogen bonds.

The concatenated insertions from Table 1 were evaluated individually to verify the AFS is a proper indication of a library member’s capsid assembly. These validations were carried out through HPLC SEC, DLS, and thermal shift assays (Figure 4). The HPLC SEC assembly assay shows that an appropriate “cut off” for assembling loop insertions is an AFS of 0.2. Most insertions with scores equal to or greater than 0.2 were observed to coelute with the WT MS2 VLP. The assembling insertions were then characterized by DLS to observe if they contained well-formed particles near the size of WT MS2. Those that are shown to be smaller than WT via DLS measurements were not observed to elute later as is observed with S37P (T=1, 17 nm).^34^ Since DLS measurements of polydisperse samples can be skewed significantly, the exact size of these VLPs cannot be determined with DLS alone and is likely an artifact from DLS. Furthermore, we observed that samples with greater radii (supplemental Figure S3) were likely beginning to aggregate out of solution, which will also greatly obscure DLS measurements. The thermal shift assay shows that while some insertions have similar melting temperatures to WT MS2, others have decreased melting temperature, and likely have lowered stability in solution compared to WT.

**Figure 4.**
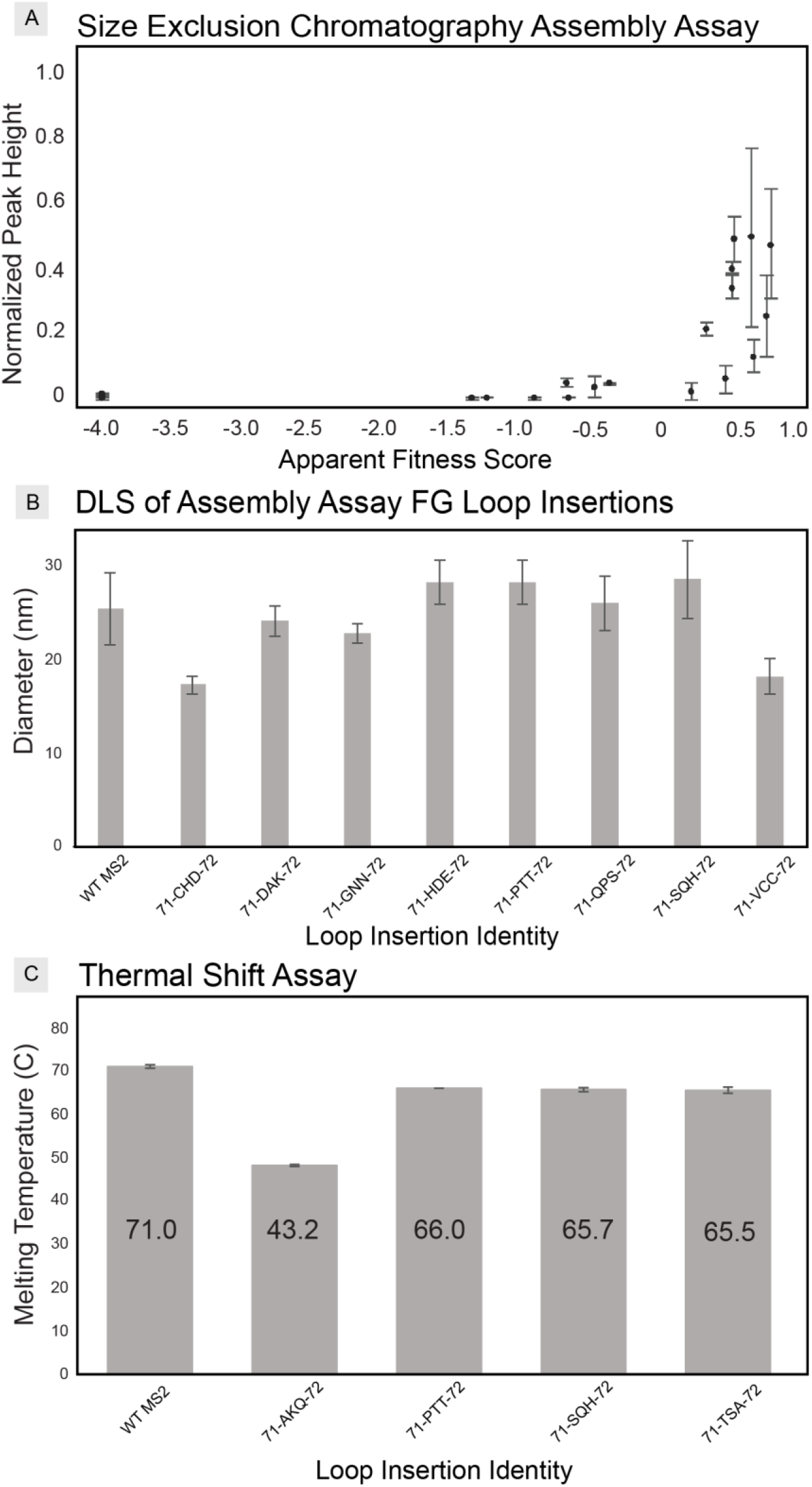
Validations of the AFL. (A) Assembly assay of the 20 concatenated peptide insertions (see Table 1). Peak heights were normalized against WT MS2 by HPLC SEC (n = 3) versus the respective AFS. (B) DLS measurements of well-formed capsids in the assembly assay show defined particle sizes in the range of the MS2 VLPs. (C) Thermal shift assay of select well-formed capsids (n = 3).

### Analyzing the Physical Property Trends of Tripeptide Insertions

To assess the physical properties tolerated at each position quantitatively, we calculated how well each of ten critical physical properties influenced assembly (Figure 5). This analysis highlights trends for each position independently and demonstrates that large polar areas, polarities, lengths, and flexibilities are favored at every position of the tripeptide insertion. Additionally, large volumes (steric bulk), nonpolar areas, and hydrophobicity are disfavored in every position.

**Figure 5.**
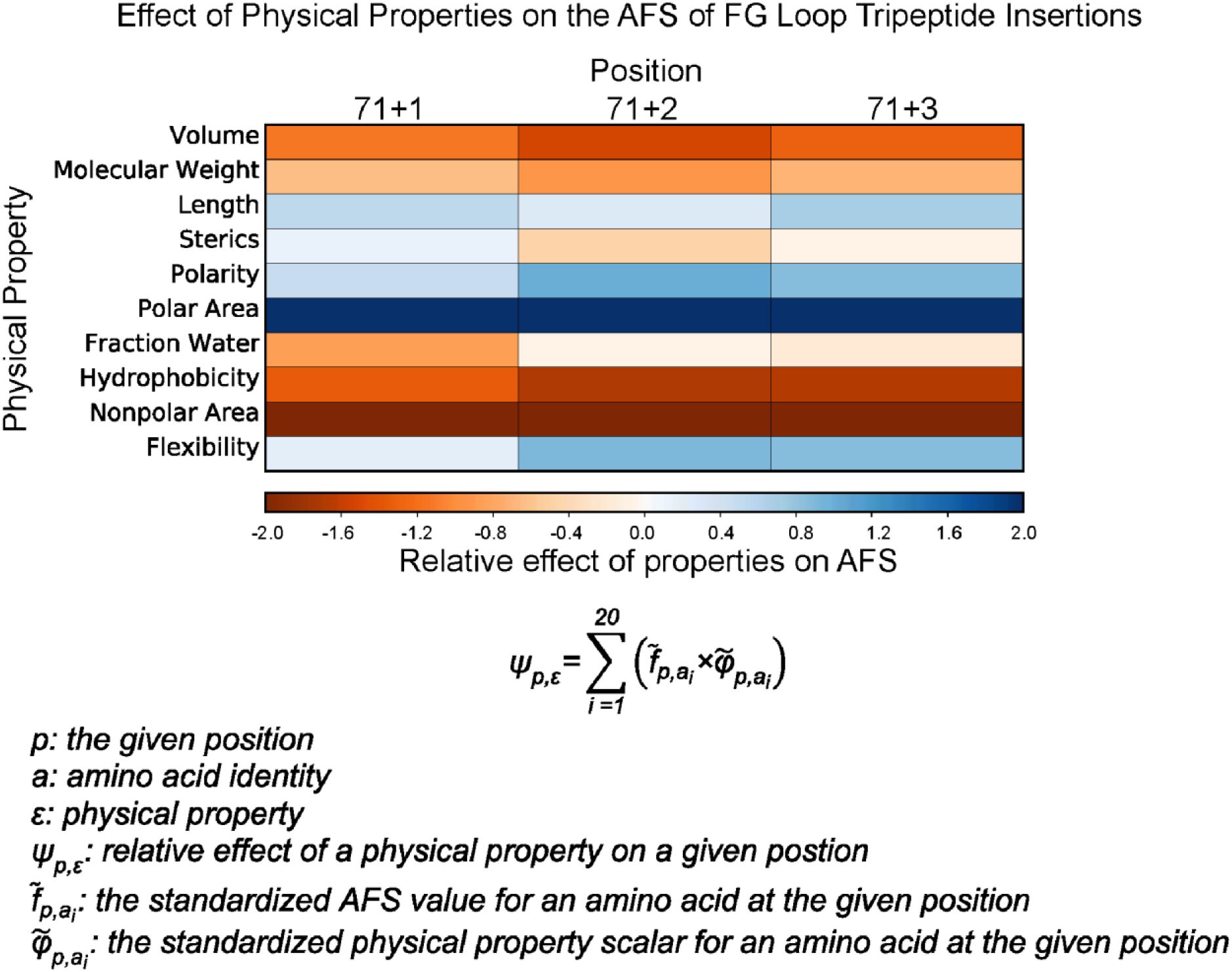
The effect of physical properties on the FG tripeptide insertion AFS. The relative effect is calculated as the sum of the products of standardized AFS values and standardized scalars of a defined physical property for the twenty amino acids in a given position. More positive scores (blue) indicate a preference for high values of the physical property, while more negative scores (red) show a bias against high values of the physical property.

This analysis identifies key differences between the positions of the FG tripeptide insertions. We observe that the first position (71+1) best tolerates steric bulk and is least favorable towards the properties of polarity, fraction of bound water (average number of bound water molecules to a residue), and flexibility. This indicates that the environment for 71+1 is distinct from that of 71+3. We hypothesize that there may be hydrophobic contacts between 71+1 and the valine-rich FG loop of adjacent monomers. These calculations highlight the differences in behavior of the positions in the tripeptide, which initially were difficult to identify by eye in the AFL.

Moving forward, these calculations can be used for engineering noncanonical amino acids in a tripeptide insertion between residues 71 and 72. For example, comparing the effects of physical properties could help predict whether an aryl azide handle or other bio-orthogonal functional groups could be a part of the FG loop tripeptide insertion and which position would best tolerate the foreign side chain. These handles could help engineer complex systems with the MS2 VLP through well-established techniques, such as copper-free click-chemistry or photo-crosslinking.^35,36^

### Uncovering Positional Dependences of Residues in Peptide Insertions

To examine positional trends, we identified variants that contained leucine, lysine, or proline in any position, and we plotted the AFS values of variants in which one of each of these residues was held constant in every position of the tripeptide insertion (Figure 6). Additionally, we used PHYRE2 modeling to predict where the inserted amino acids may be located in 3D space relative to the native residues of the FG loop. While we observed that hydrophobic and bulky residues perform poorly in these tripeptide insertions, these residues were more likely to allow for assembly when placed in position 71+1. The models of the peptide insertions suggest that the residue at position 71+1 occupies the same space as the native valine (V72). This may explain how high scoring insertions, such as FTN (AFS = 0.732) and WKT (AFS = 0.207) (assembly assessed by SEC in Supplementary Figure 5), contain hydrophobic residues in the solvent accessible pore. When examining lysine in each position (Figure 6B), the reverse trend becomes apparent. The charged residue is more tolerated in positions 71+2 and 71+3 but biased against in the first position. Modeling via PHYRE2 also suggests that proline best accommodates the hairpin turn of the loop in the middle of the tripeptide insertion, and may explain the bias for proline in position 71+2. The PHYRE2 modeling provides structural insight to trends in the AFL for peptide insertions at a residue level.

**Figure 6.**
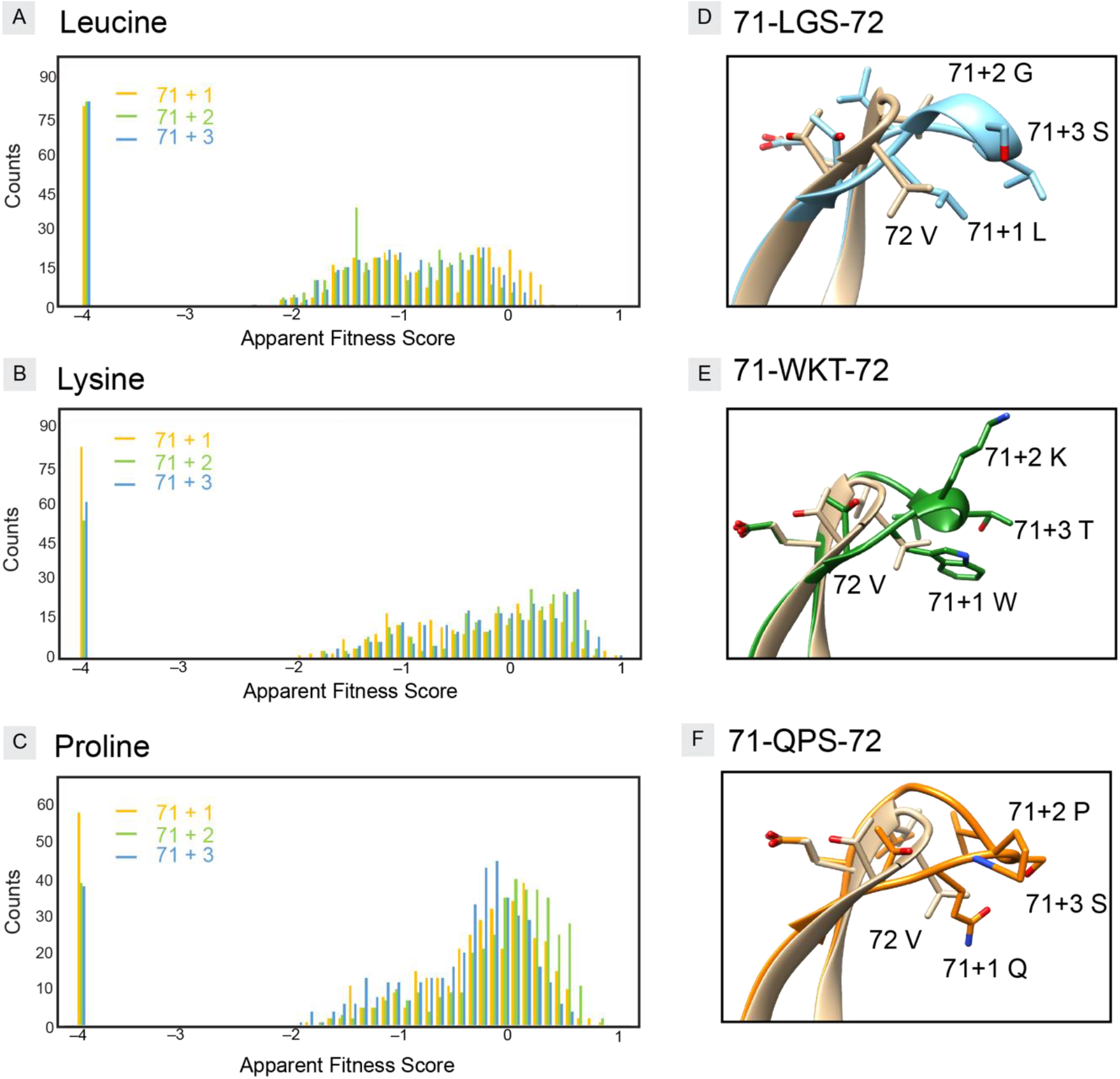
Histogram of AFS for (A) leucine, (B) lysine, or (C) proline containing peptide insertions demonstrate how the distribution of AFS is altered when restricted to a particular position of the insertion. (D, E, F) PHYRE2 modeling of assembling insertions aligned with WT (beige) structure.^37^

### Visualizing the Effects of High Localizations of Charge Affect Assembly

The FG loop insertions a show tolerance for high localizations of charge. Complementary charges tend to increase AFS for an insertion. Of the four charged residues at pH 7, the insertion library favors aspartic acid and lysine more than arginine or glutamic acid (see Table 1). Interestingly, aspartic acid performs better in peptide insertions than glutamic acid, while asparagine and glutamine perform similarly. This result could suggest that the tolerated spatial region for negative charge is more limited than is the case for the amide-containing side chains. This trend is also seen when comparing lysine and arginine: lysine is well tolerated, while arginine is disfavored.

However, when charges are balanced, the AFS of variants with multiple charged residues in their peptide insertions are higher on average. While the AFL of tripeptide insertions shows that complementary charges are favored, net charges of +/−1 also allow for VLP assembly (Figure 7). This analysis indicates that two charges in these positions are not well tolerated unless the third residue in the peptide insertion is oppositely charged. We do see great variance in AFS for each category of charges in Figure 7. This may be due to residues with near neutral pKa values easily changing their protonation states when adjacent to charged residues in the insertion. For example, the insertions, EHR (AFS = 0.540) and RHE (AFS = 0.454), suggest that the charge interactions are more favorable than the unfavorable placement of the charged residue in the first position. This is promising for a common tripeptide motif RGD; both the RGD sequence and its analogs enhance cellular specificity and uptake of proteins.^38,39^ The RGD insertion in this library is an assembling mutant (AFS = 0.378), which is consistent with high-scoring variants containing insertions with oppositely charged and flexible residues. Ultimately, the interactions of charges between residues in a peptide insertion are complex and are likely involve both electrostatic forces and hydrogen bond stabilization.

**Figure 7.**
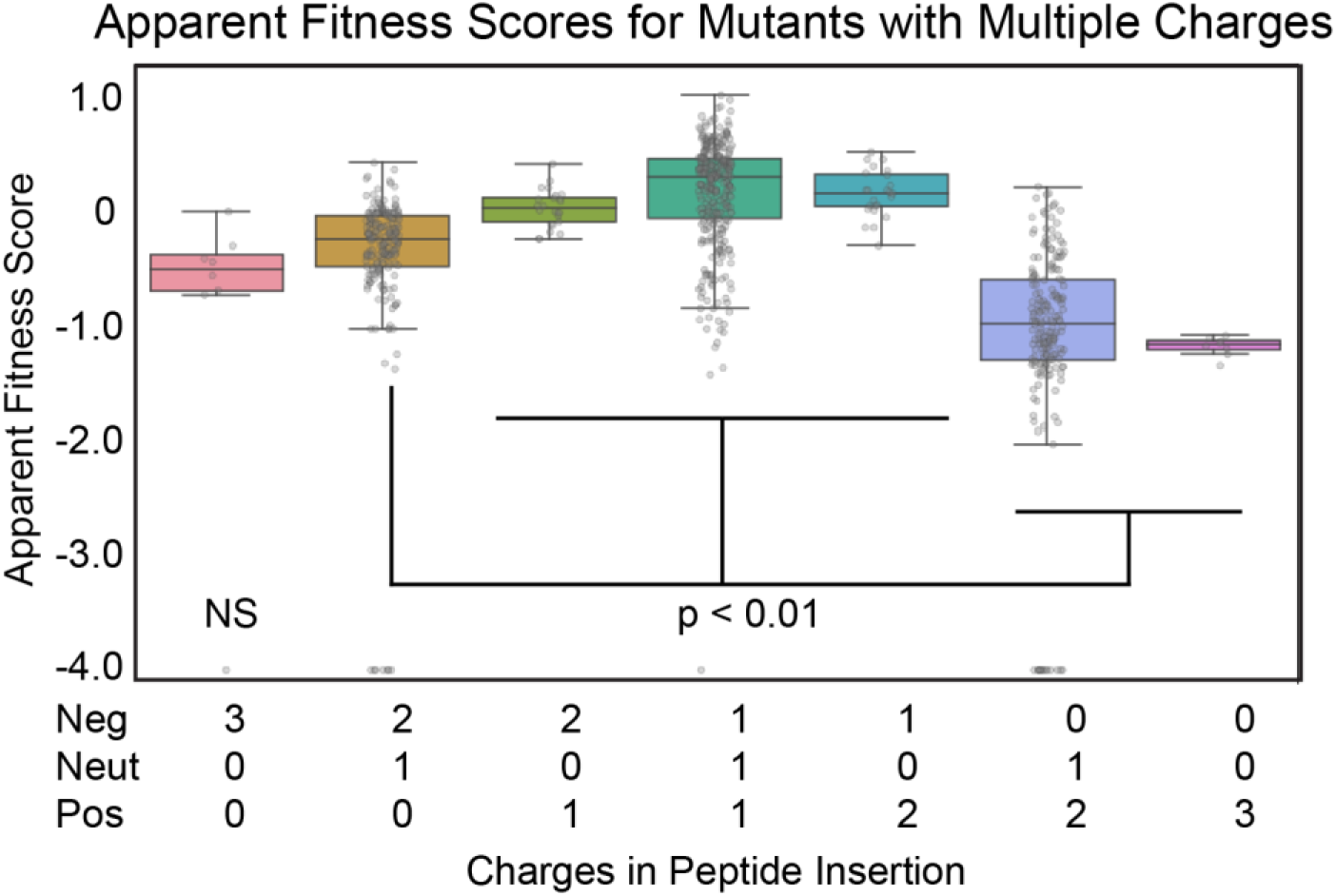
A box and whisker plot of peptide insertions with at least two charged residues. The p values shown demonstrate the separation of populations with at least two opposite charges present versus multiple identical charges. The population with negative charges is found to be not significant (NS) to all other populations of charged peptide insertions in the library.

## CONCLUSIONS

We identified a region in the MS2 CP for a large, high-quality library of peptide insertions without requiring dimer fusions to stabilize the assembled VLP. The library approach demonstrated which physical properties were allowed in the FG loop peptide insertions. The FG loop was tolerant of residues that were flexible and could hydrogen bond, while further analysis showed trends that differed for each position of the tripeptide insertion. This region of the protein shows great promise for further engineering, since it tolerates a diverse set of insertion sequences. Using the information from this tripeptide insertion library, future work includes exploring peptide insertions of increased length and employing SyMAPS to engineer the MS2 VLP further. The ability to engineer peptide insertions in a high-throughput manner will enable researchers in the field to increase the efficacy of MS2 as a drug delivery system and generate novel vaccines.

## Supporting information

Sequencing text files, python code, and primer descriptions

## ASSOCIATED CONTENT

Additional figures and supplemental information (PDF)

Primers, barcodes, and sequencing text file descriptions (XSLX)

Sequencing text files (TXT)

AFL calculations and analysis (IPNYB)

## AUTHOR INFORMATION

### Author Contributions

S.A.R., E.C.H, M.B.F, and D.T.E. conceived this project. S.A.R. and E.C.H. performed experiments for this project and led analysis of the apparent fitness landscapes. B.C.I. performed the physical properties calculations. S.A.R., E.C.H., M.B.F., and D.T.E. contributed to analyses and wrote the manuscript. All of the authors reviewed and contributed to the manuscript.

### Funding Sources

Portions of this project were funded by National Science Foundation Award CBET-1844336 to D.T.E., Army Research Office Award W911NF-16-1-0169 to D.T.E., and the BASF CARA program at the University of California, Berkeley. E.C.H. was supported under by the Department of Defense, Air Force Office of Scientific Research, National Defense Science and Engineering Graduate (NDSEG) Fellowship (32 CFR 168a).

## ACKNOWLEDGMENTS

The authors thank Daniel Brauer, Nolan Kennedy, Michael Jewett, and Eric Roth for helpful discussions and instrumentation to conduct these analyses. The sequencing was performed by the DNA Technologies and Expression Analysis Cores at the University of California Davis Genome Center, supported by National Institutes of Health Shared Instrumentation Grant S10 OD010786. This work made use of the EPIC facility of Northwestern University’s NUANCE Center, which has received support from the Soft and Hybrid Nanotechnology Experimental (SHyNE) Resource (NSF ECCS-1542205); and the MRSEC program (NSF DMR-1720139) at the Materials Research Center. This work made use of the Keck-II facility of Northwestern University’s NUANCE Center, which has received support from the Soft and Hybrid Nanotechnology Experimental (SHyNE) Resource (NSF ECCS-1542205); the MRSEC program (NSF DMR-1720139) at the Materials Research Center; the International Institute for Nanotechnology (IIN); the Keck Foundation; and the State of Illinois, through the IIN.

## ABBREVIATIONS

AFL: apparent fitness landscape
AFS: apparent fitness score
CP: coat protein
DLS: dynamic light scattering
FPLC: fast protein liquid chromatography
HPLC: high pressure liquid chromatography
SEC: size exclusion chromatography
SyMAPS: systematic mutagenesis and assembled particle selection
VLP: virus-like particle

---

SYNOPSIS Table of Contents Graphic only.

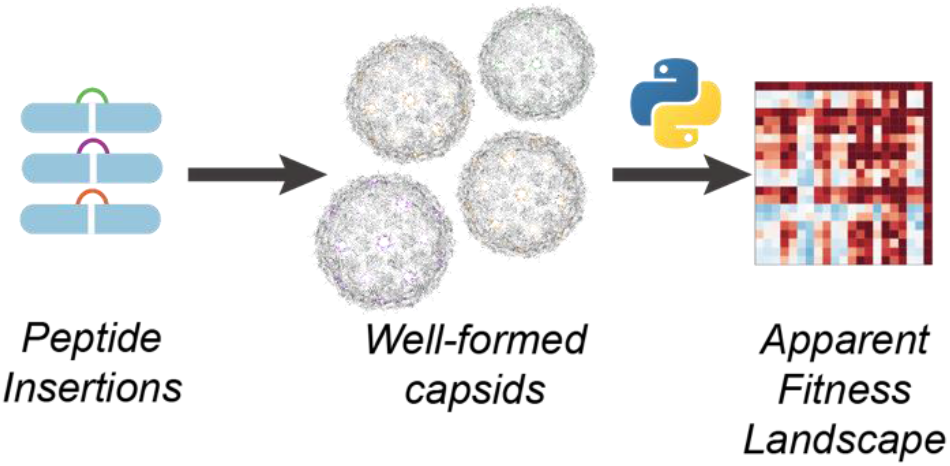

## Notes

### Competing Interest Statement

The authors have declared no competing interest.

## REFERENCES

(1) Shirbaghaee, Z.; Bolhassani, A. Different Applications of Virus-like Particles in Biology and Medicine: Vaccination and Delivery Systems. Biopolymers 2016, 105 (3), 113–132. https://doi.org/10.1002/bip.22759.

(2) Ding, X.; Liu, D.; Booth, G.; Gao, W.; Lu, Y. Virus-Like Particle Engineering: From Rational Design to Versatile Applications. Biotechnology Journal 2018, 13 (5), 1700324. https://doi.org/10.1002/biot.201700324.

(3) Golmohammadi, R.; Valegård, K.; Fridborg, K.; Liljas, L. The Refined Structure of Bacteriophage MS2 at 2·8 Å Resolution. Journal of Molecular Biology 1993, 234 (3), 620–639. https://doi.org/10.1006/jmbi.1993.1616.

(4) Plevka, P.; Tars, K.; Liljas, L. Structure and Stability of Icosahedral Particles of a Covalent Coat Protein Dimer of Bacteriophage MS2. Protein science : a publication of the Protein Society 2009, 18 (8), 1653–1661. https://doi.org/10.1002/pro.184.

(5) Lua, L. H. L.; Connors, N. K.; Sainsbury, F.; Chuan, Y. P.; Wibowo, N.; Middelberg, A. P. J. Bioengineering Virus-like Particles as Vaccines. Biotechnology and Bioengineering. March 2014, pp 425–440. https://doi.org/10.1002/bit.25159.

(6) Farkas, M. E.; Aanei, I. L.; Behrens, C. R.; Tong, G. J.; Murphy, S. T.; Neil, J. P. O.; Francis, M. B. PET Imaging and Biodistribution of Chemically Modified Bacteriophage MS2. Molecular Pharmaceutics 2013, 10, 69–76.

(7) Aanei, I. L.; Huynh, T.; Seo, Y.; Francis, M. B. Vascular Cell Adhesion Molecule-Targeted MS2 Viral Capsids for the Detection of Early-Stage Atherosclerotic Plaques. Bioconjugate Chemistry 2018, 29 (8), 2526–2530. https://doi.org/10.1021/acs.bioconjchem.8b00453.

(8) Remaut, E.; Waele, P. de; Marmenout, A.; Stanssens, P.; Fiers, W. Functional Expression of Individual Plasmid-Coded RNA Bacteriophage MS2 Genes. The EMBO Journal 1982, 1 (2), 205–209. https://doi.org/10.1002/j.1460-2075.1982.tb01148.x.

(9) Aanei, I. L.; Elsohly, A. M.; Farkas, M. E.; Netirojjanakul, C.; Regan, M.; Taylor Murphy, S.; O’Neil, J. P.; Seo, Y.; Francis, M. B. Biodistribution of Antibody-MS2 Viral Capsid Conjugates in Breast Cancer Models. Molecular Pharmaceutics 2016, 13 (11), 3764–3772. https://doi.org/10.1021/acs.molpharmaceut.6b00566.

(10) Zhang, X.; Xin, L.; Li, S.; Fang, M.; Zhang, J.; Xia, N.; Zhao, Q. Lessons Learned from Successful Human Vaccines: Delineating Key Epitopes by Dissecting the Capsid Proteins. Human Vaccines & Immunotherapeutics 2015, 11 (5), 1277–1292. https://doi.org/10.1080/21645515.2015.1016675.

(11) Smith, G. P.; Petrenko, V. A. Phage Display. Chemical Reviews 1997, 97 (2), 391–410. https://doi.org/10.1021/cr960065d.

(12) Zhai, L.; Peabody, J.; Pang, Y. Y. S.; Schiller, J.; Chackerian, B.; Tumban, E. A Novel Candidate HPV Vaccine: MS2 Phage VLP Displaying a Tandem HPV L2 Peptide Offers Similar Protection in Mice to Gardasil-9. Antiviral Research 2017, 147, 116–123. https://doi.org/10.1016/j.antiviral.2017.09.012.

(13) Caldeira, J.; Bustos, J.; Peabody, J.; Chackerian, B.; Peabody, D. S. Epitope-Specific Anti-HCG Vaccines on a Virus like Particle Platform. PLoS ONE 2015, 10 (10), 1–11. https://doi.org/10.1371/journal.pone.0141407.

(14) Tumban, E.; Muttil, P.; Escobar, C. A. A.; Peabody, J.; Wafula, D.; Peabody, D. S.; Chackerian, B. Preclinical Refinements of a Broadly Protective VLP-Based HPV Vaccine Targeting the Minor Capsid Protein, L2. Vaccine 2015, 33 (29), 3346–3353. https://doi.org/10.1016/j.vaccine.2015.05.016.

(15) Brito, C. R. N.; McKay, C. S.; Azevedo, M. A.; Santos, L. C. B.; Venuto, A. P.; Nunes, D. F.; D’Ávila, D. A.; Rodrigues da Cunha, G. M.; Almeida, I. C.; Gazzinelli, R. T.; et al. Virus-like Particle Display of the α-Gal Epitope for the Diagnostic Assessment of Chagas Disease. ACS Infectious Diseases 2016, 2 (12), 917–922. https://doi.org/10.1021/acsinfecdis.6b00114.

(16) Peabody, D. S.; Manifold-Wheeler, B.; Medford, A.; Jordan, S. K.; do Carmo Caldeira, J.; Chackerian, B. Immunogenic Display of Diverse Peptides on Virus-like Particles of RNA Phage MS2. Journal of Molecular Biology 2008, 380 (1), 252–263. https://doi.org/10.1016/j.jmb.2008.04.049.

(17) Hartman, E. C.; Jakobson, C. M.; Favor, A. H.; Lobba, M. J.; Álvarez-Benedicto, E.; Francis, M. B.; Tullman-Ercek, D. Quantitative Characterization of All Single Amino Acid Variants of a Viral Capsid-Based Drug Delivery Vehicle. Nature Communications 2018, 9 (1), 1385. https://doi.org/10.1038/s41467-018-03783-y.

(18) Brauer, D. D.; Hartman, E. C.; Bader, D. L. v.; Merz, Z. N.; Tullman-Ercek, D.; Francis, M. B. Systematic Engineering of a Protein Nanocage for High-Yield, Site-Specific Modification. Journal of the American Chemical Society 2019, 141 (9), 3875–3884. https://doi.org/10.1021/jacs.8b10734.

(19) Hartman, E. C.; Lobba, M. J.; Favor, A. H.; Robinson, S. A.; Francis, M. B.; Tullman-Ercek, D. Experimental Evaluation of Coevolution in a Self-Assembling Particle. Biochemistry 2019, 58 (11), 1527–1538. https://doi.org/10.1021/acs.biochem.8b00948.

(20) Stonehouse, N. J.; Valegård, K.; Golmohammadi, R.; van den Worm, S.; Walton, C.; Stockley, P. G.; Liljas, L. Crystal Structures of MS2 Capsids with Mutations in the Subunit FG Loop. Journal of Molecular Biology 1996, 256 (2), 330–339. https://doi.org/10.1006/jmbi.1996.0089.

(21) Koning, R. I.; Gomez-Blanco, J.; Akopjana, I.; Vargas, J.; Kazaks, A.; Tars, K.; Carazo, J. M.; Koster, A. J. Asymmetric Cryo-EM Reconstruction of Phage MS2 Reveals Genome Structure in Situ. Nature Communications 2016, 7, 12524. https://doi.org/10.1038/ncomms12524.

(22) Guzman, L. M.; Belin, D.; Carson, M. J.; Beckwith, J. Tight Regulation, Modulation, and High-Level Expression by Vectors Containing the Arabinose P(BAD) Promoter. Journal of Bacteriology 1995, 177 (14), 4121–4130. https://doi.org/10.1128/jb.177.14.4121-4130.1995.

(23) Hietpas, R. T.; Jensen, J. D.; Bolon, D. N. A. Experimental Illumination of a Fitness Landscape. Proceedings of the National Academy of Sciences of the United States of America 2011, 108 (19), 7896–7901. https://doi.org/10.1073/pnas.1016024108.

(24) Engler, C.; Gruetzner, R.; Kandzia, R.; Marillonnet, S. Golden Gate Shuffling: A One-Pot DNA Shuffling Method Based on Type IIs Restriction Enzymes. PLoS ONE 2009, 4 (5), e5553. https://doi.org/10.1371/journal.pone.0005553.

(25) Fauchère, J.-L.; Charton, M.; Kier, L. B.; Verloop, A.; Pliska, V. Amino Acid Side Chain Parameters for Correlation Studies in Biology and Pharmacology. International Journal of Peptide and Protein Research 2009, 32 (4), 269–278. https://doi.org/10.1111/j.1399-3011.1988.tb01261.x.

(26) Krigbaum, W. R.; Komoriya, A. Local Interactions as a Structure Determinant for Protein Molecules II. Biochem. et Biophys. Acta 1979, 576 (1), 204–228.

(27) Sandberg, M.; Eriksson, L.; Jonsson, J.; Sjöström, M.; Wold, S. New Chemical Descriptors Relevant for the Design of Biologically Active Peptides. A Multivariate Characterization of 87 Amino Acids. Journal of Medicinal Chemistry 1998, 41 (14), 2481–2491. https://doi.org/10.1021/jm9700575.

(28) Zimmerman, J. M.; Eliezer, N.; Simha, R. The Characterization of Amino Acid Sequences in Proteins by Statistical Methods. Journal of Theoretical Biology 1968, 21 (2), 170–201. https://doi.org/10.1016/0022-5193(68)90069-6.

(29) Vihinen, M.; Torkkila, E.; Riikonen, P. Accuracy of Protein Flexibility Predictions. Proteins: Structure, Function, and Genetics 1994, 19 (2), 141–149. https://doi.org/10.1002/prot.340190207.

(30) Fauchére, J.-L.; Charton, M.; Kier, L. B.; Verloop, A.; Pliska, V. Amino Acid Side Chain Parameters for Correlation Studies in Biology and Pharmacology. International Journal of Peptide and Protein Research 2009, 32 (4), 269–278. https://doi.org/10.1111/j.1399-3011.1988.tb01261.x.

(31) Handbook of Biochemistry and Molecular Biology; Lundblad, R., Macdonald, F., Eds.; CRC Press, 2010. https://doi.org/10.1201/b10501.

(32) Pontius, J.; Richelle, J.; Wodak, S. J. Deviations from Standard Atomic Volumes as a Quality Measure for Protein Crystal Structures. Journal of Molecular Biology 1996, 264 (1), 121–136. https://doi.org/10.1006/JMBI.1996.0628.

(33) Charton, M. Protein Folding and the Genetic Code: An Alternative Quantitative Model. Journal of Theoretical Biology 1981, 91 (1), 115–123. https://doi.org/10.1016/0022-5193(81)90377-5.

(34) Asensio, M. A.; Morella, N. M.; Jakobson, C. M.; Hartman, E. C.; Glasgow, J. E.; Sankaran, B.; Zwart, P. H.; Tullman-Ercek, D. A Selection for Assembly Reveals That a Single Amino Acid Mutant of the Bacteriophage MS2 Coat Protein Forms a Smaller Virus-like Particle. Nano Letters 2016, 16 (9), 5944–5950. https://doi.org/10.1021/acs.nanolett.6b02948.

(35) Chin, J. W.; Schultz, P. G. In Vivo Photocrosslinking with Unnatural Amino Acid Mutagenesis. ChemBioChem 2002, 3 (11), 1135–1137. https://doi.org/10.1002/1439-7633(20021104)3:11<1135::AID-CBIC1135>3.0.CO;2-M.

(36) Joiner, C. M.; Breen, M. E.; Clayton, J.; Mapp, A. K. A Bifunctional Amino Acid Enables Both Covalent Chemical Capture and Isolation of in Vivo Protein-Protein Interactions. ChemBioChem 2017, 18 (2), 181–184. https://doi.org/10.1002/cbic.201600578.

(37) Kelley, L. A.; Mezulis, S.; Yates, C. M.; Wass, M. N.; Sternberg, M. J. E. The Phyre2 Web Portal for Protein Modeling, Prediction and Analysis. Nature Protocols 2015, 10 (6), 845–858. https://doi.org/10.1038/nprot.2015.053.

(38) Danhier, F.; le Breton, A.; Préat, V. RGD-Based Strategies To Target Alpha(v) Beta(3) Integrin in Cancer Therapy and Diagnosis. Molecular Pharmaceutics 2012, 9 (11), 2961–2973. https://doi.org/10.1021/mp3002733.

(39) Mokhtarieh, A. A.; Kim, S.; Lee, Y.; Chung, B. H.; Lee, M. K. Novel Cell Penetrating Peptides with Multiple Motifs Composed of RGD and Its Analogs. Biochemical and Biophysical Research Communications 2013, 432 (2), 359–364. https://doi.org/10.1016/j.bbrc.2013.01.096.

