## Supplemental figures for "Engineering a virus-like particle to display peptide insertions using an apparent fitness landscape"

**Figure S1.** The AFL grouped by identity at position 1 (71+1)

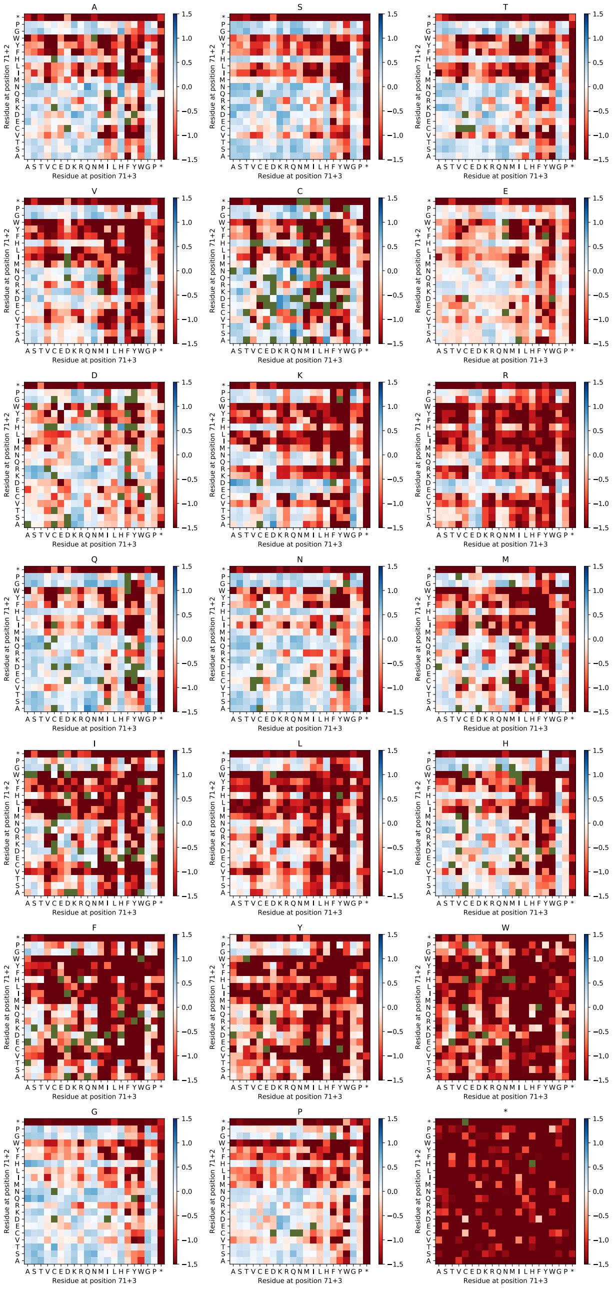

**Figure S2.** The AFL grouped by identity at position 3 (71+3)

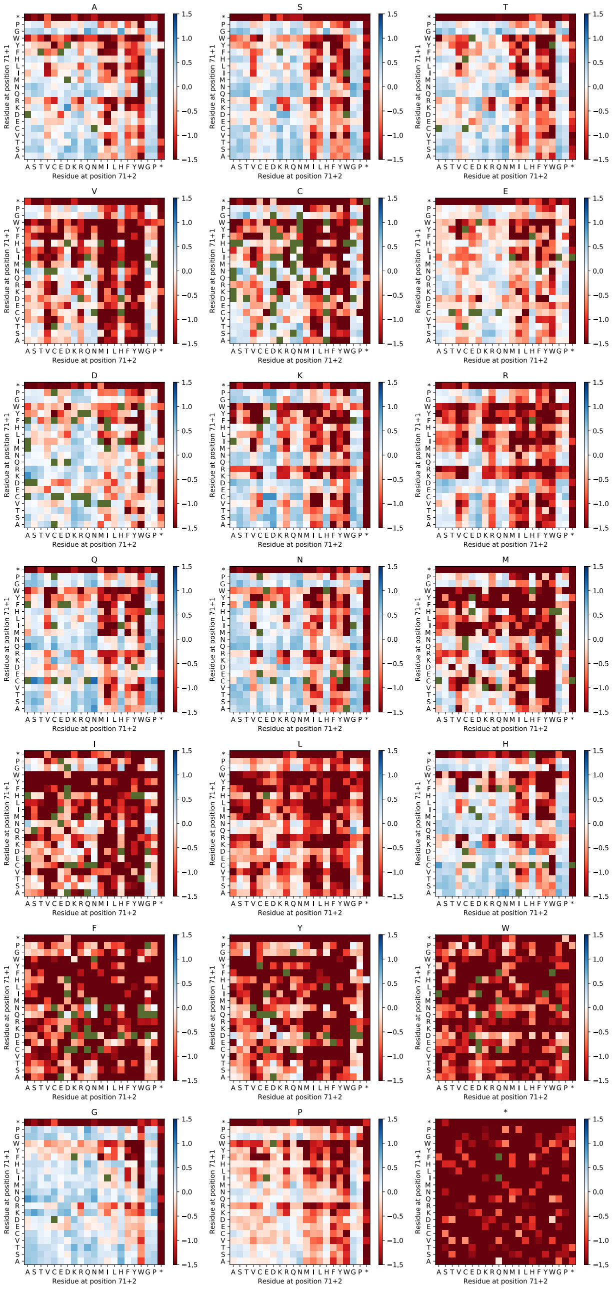

**Figure S3.** Dynamic Light Scattering measurements of assembly assay insertions show distinct radii for VLPs

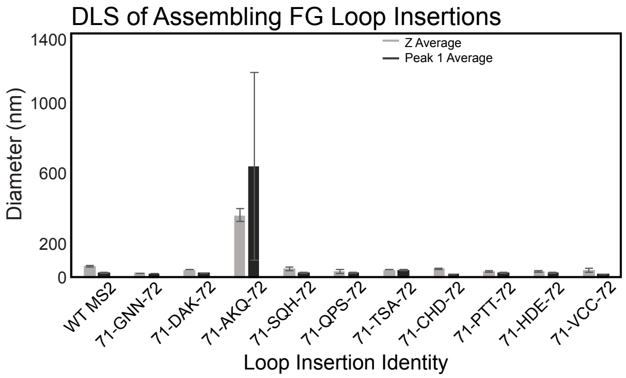


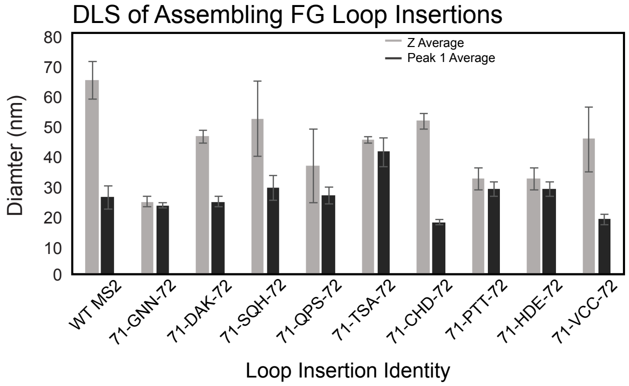

**Figure S4.** Transmission Electron Micrographs of (A) WT MS2, (B) 71-PTT-72, and (C) 71-DAK-72

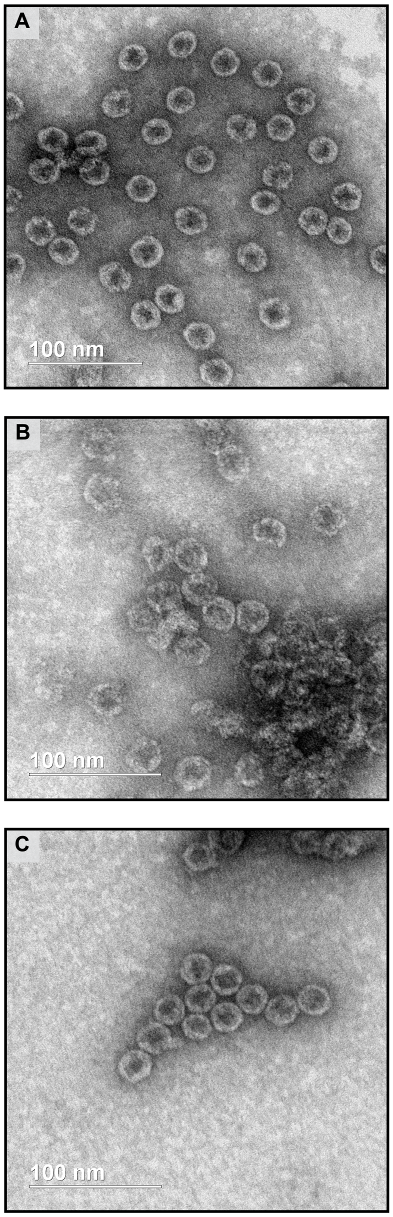

**Figure S5.** (A) HPLC SEC chromatograms show characteristic capsid peak between 7.3-9.3 min. (B) DLS measurements show the diameter of retrievable FG loop insertions have a diameter roughly 20-30 nm similar to WT MS2 (27 nm).

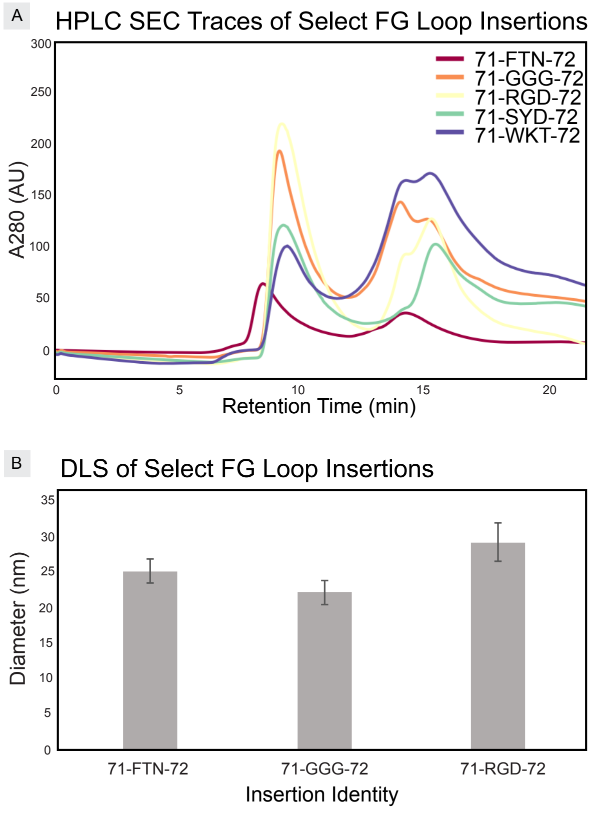
