## Supplementary material for "Engineering a virus-like particle to display peptide insertions using an apparent fitness landscape": Sequencing text files, python code, and primer descriptions: SupplementalFile_SequencingAnalysis.ipynb

```
# %% [code]
'''This notebook is walks through the data processesing of txt files for generating the AFL from sequences already undergone quality control and trimming'''

# %% [code]
from mpl_toolkits.mplot3d import Axes3D
from Bio.Seq import Seq
from Bio import SeqIO
from Bio.Alphabet import IUPAC
import csv as csv
import matplotlib as matplotlib
import matplotlib.pyplot as plt
import numpy as np
from numpy import inf
import math
import matplotlib.patches as mpatches
import numpy.ma as ma
from matplotlib import cm
from matplotlib.colors import ListedColormap, LinearSegmentedColormap
from collections import OrderedDict

%matplotlib inline

'Note - install biopython if not already installed by running the following:' 
'pip install biopython'

# %% [code]
'''Data processing for Apparent Fitness Landscape'''

def parse_from_txt_fgins(file):
    '''This takes the aligned, merged file from terminal analysis. We first get rid of any genes that do not
    start with ATG and end with TAA. Then we append the resulting genes to a list.'''
    gene = ""
    text_file = open(file, "r") #open and read the text file containing aligned library reads
    lines = text_file.readlines()
    text_file.close()
    
    record = [] #this is list of genes that will be counted later
    for x in lines: #separate out each read to be analyzed individually
        '''The start codon follow a tab, and the stop codon should preceed a tab'''
        for i in range(100): #scan the first 100 letters for \tATG
            if x[i:i+4] == "GACT":
                record.append(x[i+1:i+16]) #append to a list for further analysis
    return record

# %% [code]
def count_codons_fgins(array_of_abundances):
    '''This function will count the abundance of every codon in each position of the peptide insertion'''
    intensities = np.zeros((64, 64, 64))

    codon_to_num = {'TTT': 0, 'TTG': 1, 'TTA': 2, 'TTC': 3, 
                         'CTT': 4, 'CTG': 5, 'CTA': 6, 'CTC': 7,
                         'ATT': 8, 'ATG': 9, 'ATA': 10, 'ATC': 11,
                         'GTT': 12, 'GTG': 13, 'GTA': 14, 'GTC': 15,
                         'TAT': 16, 'TAG': 17, 'TAA': 18, 'TAC': 19,
                         'CAT': 20, 'CAG': 21, 'CAA': 22, 'CAC': 23,
                         'AAT': 24, 'AAG': 25, 'AAA': 26, 'AAC': 27,
                         'GAT': 28, 'GAG': 29, 'GAA': 30, 'GAC': 31,
                         'TCT': 32, 'TCG': 33, 'TCA': 34, 'TCC': 35,
                         'CCT': 36, 'CCG': 37, 'CCA': 38, 'CCC': 39,
                         'ACT': 40, 'ACG': 41, 'ACA': 42, 'ACC': 43,
                         'GCT': 44, 'GCG': 45, 'GCA': 46, 'GCC': 47,
                         'TGT': 48, 'TGG': 49, 'TGA': 50, 'TGC': 51,
                         'CGT': 52, 'CGG': 53, 'CGA': 54, 'CGC': 55,
                         'AGT': 56, 'AGG': 57, 'AGA': 58, 'AGC': 59,
                         'GGT': 60, 'GGG': 61, 'GGA': 62, 'GGC' :63}

    for dna_read in array_of_abundances: #dna_read counts where you are in sequencing reads
        if dna_read[0:3] == 'ACT' and dna_read[12:15] == 'GTT':
            codon_one = codon_to_num[dna_read[3:6]]
            codon_two = codon_to_num[dna_read[6:9]]
            codon_three = codon_to_num[dna_read[9:12]]
            
            intensities[codon_one,codon_two,codon_three] += 1
    return intensities

# %% [code]
def translate_NNK_array_3d(nparray1):
    '''Takes the codon-counted np array and turns it into a amino acid np array, discarding 
    anyt codon that ends in C or A and is thus not from an NNK library'''
    
    AA_code = {44: 0, 45: 0, 46: 0, 47: 0, 32: 1, 33: 1, 34: 1, 35: 1, 56: 1, 59: 1, 
                 40: 2, 41: 2, 42: 2, 43: 2, 12: 3, 13: 3, 14: 3, 15: 3, 48: 4, 51: 4, 
                 29: 5, 30: 5, 28: 6, 31: 6, 25: 7, 26: 7, 52: 8, 53: 8, 54: 8, 55: 8, 
                 57: 8, 58: 8, 21: 9, 22: 9, 24: 10, 27: 10, 9: 11, 8: 12, 10: 12, 11: 12, 
                 1: 13, 2: 13, 4: 13, 5: 13, 6: 13, 7: 13, 20: 14, 23: 14, 0: 15, 3: 15, 
                 16: 16, 19: 16, 49: 17, 60: 18, 61: 18, 62: 18, 63: 18, 36: 19, 37: 19, 
                 38: 19, 39: 19, 17: 20, 18: 20, 50: 20}
    
    '''Rearranged so output is in a logical order'''
    
    AAs = ['A', 'S', 'T', 'V', 'C', 'E', 'D', 'K', 'R', 'Q', 'N', 'M', 'I', 'L', 
           'H', 'F', 'Y', 'W', 'G', 'P', '*']

    
    NNK_only = [0, 1, 4, 5, 8, 9, 12, 13, 16, 17, 20, 21,
                    24, 25, 28, 29, 32, 33, 36, 37, 40, 41,
                    44, 45, 48, 49, 52, 53, 56, 57, 60, 61]
    
    array_translated = np.zeros((21, 21, 21))
    
    for n in NNK_only:
        for i in NNK_only:
            for w in NNK_only:#Loop through NNK_only (which is codon numbers excluding those ending in C or A)
                x = AA_code[n] #Get the amino acid location where the codon row should be added
                y = AA_code[i]
                z = AA_code[w]
                array_translated[x,y,z] += nparray1[n,i,w] #Add the entire row to the appropriate amino acid row
                
    return array_translated

# %% [code]
def fgins_processing(text_file):
    '''A distinct function for processesing, counting, and translating sequences for the FG loop peptide insertion'''
    parsed_text_file = parse_from_txt_fgins(text_file)
    sliced_list_of_dnas = count_codons_fgins(parsed_text_file)
    translated_3d_array = translate_NNK_array_3d(sliced_list_of_dnas)
    return translated_3d_array

# %% [code]
def average_and_compare_arrays(array1, array2, array3, array4, array5, array6):
    '''This will generate relative abundances of amino acid sequences in the plasmid library vs teh VLP library.
    Note that array1 - array3 represent the plasmid libraries, while array4 - array6 represent the VLP library. 
    For example, replicate 1 corresponds to array1 (plasmid library) and array4 (VLP library).'''
    norm_array1 = array1/array1.sum()
    norm_array2 = array2/array2.sum()
    norm_array3 = array3/array3.sum()
    norm_array4 = array4/array4.sum()
    norm_array5 = array5/array5.sum()
    norm_array6 = array6/array6.sum()
    
    rep1_fgins = np.divide(norm_array4, norm_array1)
    rep2_fgins = np.divide(norm_array5, norm_array2)
    rep3_fgins = np.divide(norm_array6, norm_array3)

    array_of_replicates = np.array([rep1_fgins, rep2_fgins, rep3_fgins]) 
    
    #this array of arrays means we can use axis=0 to get the mean
    
    mean_array_fgins = np.nanmean(array_of_replicates, axis=0)
    
    for i in range(21):
        for n in range(21):
            for m in range(21):
                mean_array_fgins[i,n,m] = max(mean_array_fgins[i,n,m], .0001)
    '''The apparent fitness score is then determined by a log10 scale'''
    log_mean_fgins = np.log10(mean_array_fgins)
    
    return log_mean_fgins

# %% [code]
def plot_array_of_arrays(relative_percent_abundance_array, filename):
    '''Setting up function to plotting the Apparent Fitness Landscape'''
    AAs = ['A', 'S', 'T', 'V', 'C', 'E', 'D', 'K', 'R', 'Q', 'N', 'M', 'I', 'L', 
               'H', 'F', 'Y', 'W', 'G', 'P', '*']
    y_pos = np.arange(len(AAs))

    fig, ax = plt.subplots(
        figsize = (13,27)
    )
    
    data_to_plot = np.ma.masked_invalid(relative_percent_abundance_array)
    
    # to make the RedBlue Color Maps
    top = cm.get_cmap('Reds_r', 128)
    bottom = cm.get_cmap('Blues', 128)
    newcolors = np.vstack((top(np.linspace(0, 1, 128)),bottom(np.linspace(0, 1, 128))))
    newcmp = ListedColormap(newcolors, name='RedBlue') 
    newcmp.set_bad('darkolivegreen', 1)
    
    
    
    for i in range(21):
        plt.subplot(7, 3, i+1)
        im = plt.pcolormesh(data_to_plot[:,:,i], cmap=newcmp, vmax=1.5, vmin=-1.5)
        plt.colorbar(im, aspect=40)
        
        #change x and y labels if changing which position you're grouping by
        plt.ylabel("Residue at position 71+1") 
        plt.xlabel("Residue at position 71+2")
        plt.yticks(y_pos+0.5, AAs)
        plt.xticks(y_pos+0.5, AAs)
        plt.title(AAs[i])
        plt.ylim(0,21)
        plt.xlim(0,21)
    
    fig.tight_layout()
    
    plt.savefig(filename, dpi=1000, format='pdf')
    
    return print(data_to_plot.shape)

# %% [code]
def plot_array_of_arrays(relative_percent_abundance_array, filename, position_grouping):
    '''Setting up function to plotting the Apparent Fitness Landscape'''
    AAs = ['A', 'S', 'T', 'V', 'C', 'E', 'D', 'K', 'R', 'Q', 'N', 'M', 'I', 'L', 
               'H', 'F', 'Y', 'W', 'G', 'P', '*']
    y_pos = np.arange(len(AAs))

    fig, ax = plt.subplots(
        figsize = (13,27)
    )
    
    data_to_plot = np.ma.masked_invalid(relative_percent_abundance_array)
    
    # to make the RedBlue Color Maps
    top = cm.get_cmap('Reds_r', 128)
    bottom = cm.get_cmap('Blues', 128)
    newcolors = np.vstack((top(np.linspace(0, 1, 128)),bottom(np.linspace(0, 1, 128))))
    newcmp = ListedColormap(newcolors, name='RedBlue') 
    newcmp.set_bad('darkolivegreen', 1)
    
   #plot data according to grouping by specified position 
    for i in range(21):
        plt.subplot(7, 3, i+1)
        
        if position_grouping == 1: 
            im = plt.pcolormesh(data_to_plot[i,:,:], cmap=newcmp, vmax=1.5, vmin=-1.5)
            plt.ylabel("Residue at position 71+2") 
            plt.xlabel("Residue at position 71+3")
        elif position_grouping == 2:
            im = plt.pcolormesh(data_to_plot[:,i,:], cmap=newcmp, vmax=1.5, vmin=-1.5)
            plt.ylabel("Residue at position 71+1") 
            plt.xlabel("Residue at position 71+3")
        elif position_grouping == 3:
            im = plt.pcolormesh(data_to_plot[:,:,i], cmap=newcmp, vmax=1.5, vmin=-1.5)
            plt.ylabel("Residue at position 71+1") 
            plt.xlabel("Residue at position 71+2")

        
        plt.colorbar(im, aspect=40)
        plt.yticks(y_pos+0.5, AAs)
        plt.xticks(y_pos+0.5, AAs)
        plt.title(AAs[i])
        plt.ylim(0,21)
        plt.xlim(0,21)
    
        
    fig.tight_layout()
    
    plt.savefig(filename, dpi=1000, format='pdf')
    
    return print(data_to_plot.shape)

# %% [code]
def compare_and_plot_fgins(array1_text, array2_text, array3_text, array4_text, array5_text, array6_text, name, position_grouping):
    '''Combine functions from before to process sequences, calculate AFS, and return the AFL'''
    array1 = fgins_processing(array1_text)
    array2 = fgins_processing(array2_text)
    array3 = fgins_processing(array3_text)
    array4 = fgins_processing(array4_text)
    array5 = fgins_processing(array5_text)
    array6 = fgins_processing(array6_text)
    
    compared_array = average_and_compare_arrays(array1, array2, array3, array4, array5, array6)
    plot_array_of_arrays(compared_array, name, position_grouping)
    
    return compared_array

# %% [code]
#Now generate and plot the FG Peptide Insertion AFL
fgins_afl = compare_and_plot_fgins('eHTS065S24_s1_pe', 'eHTS066S25_s1_pe', 'eHTS067S26_s1_pe', 
                                   'eHTS068S27_s1_pe', 'eHTS069S28_s1_pe', 'eHTS070S29_s1_pe', 
                                   'FGPeptideInsertion_AFL_groupedpos2.pdf', 2)

# %% [code]
#For future analysis and not having to repeat AFL generation, save the AFL as txt or csv. 
np.savetxt('FGPeptideInsertion_AFL', fgins_afl.flat)

# %% [code]
#Physical Properties Calculations 

# %% [code]
def count_codons_fgins_pp(array_of_abundances):
    
    intensities = np.zeros((64, 64, 64))
    intensities_physicalProperties =  np.zeros((3,64)) #this is for creating a 3 residue length array for physical property calculations

    codon_to_num = {'TTT': 0, 'TTG': 1, 'TTA': 2, 'TTC': 3, 
                         'CTT': 4, 'CTG': 5, 'CTA': 6, 'CTC': 7,
                         'ATT': 8, 'ATG': 9, 'ATA': 10, 'ATC': 11,
                         'GTT': 12, 'GTG': 13, 'GTA': 14, 'GTC': 15,
                         'TAT': 16, 'TAG': 17, 'TAA': 18, 'TAC': 19,
                         'CAT': 20, 'CAG': 21, 'CAA': 22, 'CAC': 23,
                         'AAT': 24, 'AAG': 25, 'AAA': 26, 'AAC': 27,
                         'GAT': 28, 'GAG': 29, 'GAA': 30, 'GAC': 31,
                         'TCT': 32, 'TCG': 33, 'TCA': 34, 'TCC': 35,
                         'CCT': 36, 'CCG': 37, 'CCA': 38, 'CCC': 39,
                         'ACT': 40, 'ACG': 41, 'ACA': 42, 'ACC': 43,
                         'GCT': 44, 'GCG': 45, 'GCA': 46, 'GCC': 47,
                         'TGT': 48, 'TGG': 49, 'TGA': 50, 'TGC': 51,
                         'CGT': 52, 'CGG': 53, 'CGA': 54, 'CGC': 55,
                         'AGT': 56, 'AGG': 57, 'AGA': 58, 'AGC': 59,
                         'GGT': 60, 'GGG': 61, 'GGA': 62, 'GGC' :63}

    for dna_read in array_of_abundances: #dna_read counts where you are in sequencing reads
          if dna_read[0:3] == 'ACT' and dna_read[12:15] == 'GTT':
            
            codon_one = codon_to_num[dna_read[3:6]]
            intensities_physicalProperties[0,codon_one] += 1
            
            codon_two = codon_to_num[dna_read[6:9]]
            intensities_physicalProperties[1,codon_two] += 1
            
            codon_three = codon_to_num[dna_read[9:12]]
            intensities_physicalProperties[2,codon_three] += 1
            
            
            intensities[codon_one,codon_two,codon_three] += 1
    
    return intensities, intensities_physicalProperties


# %% [code]
'''Data processing for Apparent Fitness Landscape that I want to use for physical property calculations'''

def translate_NNK_array_pp(nparray1):
    '''Takes the codon-counted np array and turns it into a amino acid np array, discarding 
    anything that ends in C or A and is thus not from an NNK library'''
    
    AA_code = {44: 0, 45: 0, 46: 0, 47: 0, 32: 1, 33: 1, 34: 1, 35: 1, 56: 1, 59: 1, 
                 40: 2, 41: 2, 42: 2, 43: 2, 12: 3, 13: 3, 14: 3, 15: 3, 48: 4, 51: 4, 
                 29: 5, 30: 5, 28: 6, 31: 6, 25: 7, 26: 7, 52: 8, 53: 8, 54: 8, 55: 8, 
                 57: 8, 58: 8, 21: 9, 22: 9, 24: 10, 27: 10, 9: 11, 8: 12, 10: 12, 11: 12, 
                 1: 13, 2: 13, 4: 13, 5: 13, 6: 13, 7: 13, 20: 14, 23: 14, 0: 15, 3: 15, 
                 16: 16, 19: 16, 49: 17, 60: 18, 61: 18, 62: 18, 63: 18, 36: 19, 37: 19, 
                 38: 19, 39: 19, 17: 20, 18: 20, 50: 20}
    
    '''Rearranged so output is in a logical order'''
    
    AAs = ['A', 'S', 'T', 'V', 'C', 'E', 'D', 'K', 'R', 'Q', 'N', 'M', 'I', 'L', 
           'H', 'F', 'Y', 'W', 'G', 'P', '*']

    
    NNK_only = [0, 1, 4, 5, 8, 9, 12, 13, 16, 17, 20, 21,
                    24, 25, 28, 29, 32, 33, 36, 37, 40, 41,
                    44, 45, 48, 49, 52, 53, 56, 57, 60, 61]
    
    array_translate = np.zeros((3, 21))
    
    for i in NNK_only: #Loop through NNK_only (which is codon numbers excluding those ending in C or A)
        x = AA_code[i] #Get the amino acid location where the codon row should be added
        array_translate[:,x] += nparray1[:,i] #Add the entire row to the appropriate amino acid row

    return array_translate


# %% [code]
def average_and_compare_arrays_pp(array1, array2, array3, array4, array5, array6):
    
    norm_array1 = array1/array1.sum()
    norm_array2 = array2/array2.sum()
    norm_array3 = array3/array3.sum()
    norm_array4 = array4/array4.sum()
    norm_array5 = array5/array5.sum()
    norm_array6 = array6/array6.sum()
    
    rep1_ins = np.divide(norm_array4, norm_array1)
    rep2_ins = np.divide(norm_array5, norm_array2)
    rep3_ins = np.divide(norm_array6, norm_array3)

    rep1_ins [rep1_ins == inf] = np.nan 
    rep2_ins [rep2_ins == inf] = np.nan 
    rep3_ins [rep3_ins == inf] = np.nan 
    array_of_replicates = np.array([rep1_ins, rep2_ins, rep3_ins]) 
    
    #this array of arrays means we can use axis=0 to get the mean
    
    mean_array_ins = np.nanmean(array_of_replicates, axis=0)
    
    mean_array_ins[mean_array_ins == 0] = .0001
    
    log_mean_ins = np.log10(mean_array_ins)

   
    return log_mean_ins

# %% [code]
def fgins_processing_pp(text_file):
    parsed_text_file = parse_from_txt_fgins(text_file)   
    sliced_list_of_dnas, sliced_list_of_dnas_pp = count_codons_fgins_pp(parsed_text_file)
    translated_array = translate_NNK_array_pp(sliced_list_of_dnas_pp)
    
    #print(sliced_list_of_dnas_pp.shape)
    #print(parsed_text_file)
    print(translated_array.shape)
    
    return translated_array


# %% [code]
def compare_fgins_pp(array1_text, array2_text, array3_text, array4_text, array5_text, array6_text):
    array1 = fgins_processing_pp(array1_text)
    array2 = fgins_processing_pp(array2_text)
    array3 = fgins_processing_pp(array3_text)
    array4 = fgins_processing_pp(array4_text)
    array5 = fgins_processing_pp(array5_text)
    array6 = fgins_processing_pp(array6_text)
    
    compared_array = average_and_compare_arrays_pp(array1, array2, array3, array4, array5, array6)
       
    return compared_array


# %% [code]
'''Array using for physical property calculations'''
fg_pos_heatmap_pp = compare_fgins_pp('eHTS065S24_s1_pe', 'eHTS066S25_s1_pe', 'eHTS067S26_s1_pe', 
                                   'eHTS068S27_s1_pe', 'eHTS069S28_s1_pe', 'eHTS070S29_s1_pe')

# %% [code]
'''Setting up the backbones for physical property calculations.'''
#zeta physical property vector values from Andrew Favor's Thesis
zeta_posCharge = np.array([[0],[0],[0],[0],[0],[0],[0],[1],[1],[0],
                            [0],[0],[0],[0],[1],[0],[0],[0],[0],[0]])
zeta_negCharge = np.array([[0],[0],[0],[0],[0],[1],[1],[0],[0],[0],
                            [0],[0],[0],[0],[1],[0],[0],[0],[0],[0]])
zeta_volume = np.array([[91.5],[102],[126],[138.4],[114.4],[154.6],[135.2],[162.5],[196.1],[156.4],
                         [138.3],[165.9],[162.6],[163.4],[163.2],[198.8],[237.2],[209.8],[67.5],[123.4]])
zeta_molWeight = np.array([[89.09],[105.09],[119.12],[117.15],[121.15],[147.13],[133.1],[146.19],
                     [174.2],[146.15],[132.12],[149.21],[131.17],[131.17],[155.16],
                     [165.19],[181.19],[204.24],[75.07],[115.13]])
zeta_length = np.array([[2.87],[3.97],[4.11],[4.11],[4.47],[5.97],[4.74],[6.89],[7.82],[6.11],[4.58],[6.36],[4.92],
                     [4.92],[5.23],[4.62],[4.73],[7.68],[2.06],[4.11]])
zeta_sterics= np.array([[0.52],[0.53],[0.5],[0.76],[0.62],[0.68],[0.76],[0.68],[0.68],[0.68],[0.76],[0.78],[1.02],
                     [0.98],[0.7],[0.7],[0.7],[0.7],[0],[0.36]])
zeta_polarity= np.array([[0],[1.67],[1.66],[0.13],[1.48],[49.9],[49.7],[49.5],[52],[3.53],[3.38],[1.43],[0.13],
                     [0.13],[51.6],[0.35],[1.61],[2.1],[0],[1.58]])
zeta_polarArea= np.array([[76],[91],[91],[70],[74],[121],[118],[96],[132],[126],[121],[71],[71],
                     [70],[98],[73],[95],[85],[79],[60]])
zeta_fracWater= np.array([[0.28],[0.27],[0.26],[0.22],[0.11],[0.37],[0.33],[0.59],[0.34],[0.39],[0.31],[0.08],[0.12],
                     [0.16],[0.23],[0.1],[0.25],[0.15],[0.28],[0.46]])
zeta_hydrophobicity= np.array([[0.25],[-0.26],[-0.18],[0.54],[0.04],[-0.62],[-0.72],[-1.1],[-1.76],[-0.69],[-0.64],
                                [0.26],[0.73],[0.53],[-0.4],[0.61],[0.02],[0.37],[0.16],[-0.07]])
zeta_nonpolarArea= np.array([[58],[47],[69],[98],[76],[59],[41],[108],[90],[58],[39],[125],[118],
                     [118],[98],[140],[123],[163],[33],[100]])
zeta_flexibility= np.array([[0.984],[1.046],[0.997],[0.931],[0.906],[1.094],[1.069],[1.102],[1.008],[1.037],
                             [1.048],[0.952],[0.927],[0.935],[0.95],[0.915],[0.929],[0.904],[1.031],[1.049]])

#Calculating standard fitness score array
afs_meanAtResidue = np.zeros((3,1)) #setting up array for average of AFS for a specific residue, 94 is the length of the protein
afs_stdAtResidue = np.zeros((3,1))#setting up array for standard deviation of AFS for a specific residue

#before Performing statistics on a residue, need to ignore nan values
afs_meanAtResidue =  np.nanmean(fg_pos_heatmap_pp, axis=1) #nanmean masks ignores nan values
afs_stdAtResidue = np.nanstd(fg_pos_heatmap_pp, axis=1)

#Now creating standardized fitness array

standardized_Fitness_Array = np.zeros((3,21))

for i in range(3): #iterating through every residue
    for n in range(21): #iterating through every amino acid plus amber stop codon
        standardized_Fitness_Array[i,n]= (fg_pos_heatmap_pp[i,n]- afs_meanAtResidue[i])/afs_stdAtResidue[i]
        

#now need to create stanrdized physical property indices
def scale_physical_property(physical_property):
    mean_p = physical_property.mean()
    std_p = physical_property.std()
        
    scaled_0 = (physical_property - mean_p)/std_p   
    min_value = scaled_0.min()
    scaled_1 = scaled_0 - min_value
    max_value = scaled_1.max()
    scaled_property = scaled_1 / max_value
    
    scaled_propertyT = scaled_property.T #transposing 1X20 Vector to 20X1
          
    return scaled_propertyT


scaled_volume = scale_physical_property(zeta_volume)
scaled_molWeight = scale_physical_property(zeta_molWeight)
scaled_length = scale_physical_property(zeta_length)
scaled_sterics = scale_physical_property(zeta_sterics)
scaled_polarity = scale_physical_property(zeta_polarity)
scaled_polarArea = scale_physical_property(zeta_polarArea)
scaled_fractionWater = scale_physical_property(zeta_fracWater)
scaled_hydrophobicity = scale_physical_property(zeta_hydrophobicity)
scaled_nonPolarArea = scale_physical_property(zeta_nonpolarArea)
scaled_flexibility = scale_physical_property(zeta_flexibility)

phi_physicalProperties = np.concatenate((scaled_volume,scaled_molWeight,
                                        scaled_length,scaled_sterics,scaled_polarity,scaled_polarArea,
                                        scaled_fractionWater,scaled_hydrophobicity,scaled_nonPolarArea,
                                        scaled_flexibility),axis=0)

transposed_phi_physicalProperties = phi_physicalProperties.T

# Determination of Physical Property preferences
standardized_Fitness_Array_square = standardized_Fitness_Array[:,1:21] # to be compatible with matrix multiplication
'''The nan.sum fix for using dot prodcut with arrays that have nan values'''
fgpos_pp_preferences = np.where(np.isnan(standardized_Fitness_Array_square),0,standardized_Fitness_Array_square).dot(np.where(np.isnan(transposed_phi_physicalProperties),
                                                                                                       0,transposed_phi_physicalProperties))


# %% [code]
def fgpos_plot_physical_properties(physical_array, name):
    Properties = ['Volume', 'Molecular Weight', 'Length', 'Sterics', 'Polarity', 'Polar Area', 'Fraction Water', 'Hydrophobicity',
                  'Nonpolar Area', 'Flexibility']
    ytix = []
    
    for i in range(1,4):
        ytix.append(i)

    fig, ax = plt.subplots(figsize = (7,14))

    
    data_to_plot = np.ma.masked_invalid(physical_array)
   
    
    
    plt.subplot(1, 1, 1)
    ax.patch.set(hatch='', color="#7B9569")
    top = cm.get_cmap('Oranges_r', 128)
    bottom = cm.get_cmap('Blues', 128)
    newcolors = np.vstack((top(np.linspace(0, 1, 128)),
                       bottom(np.linspace(0, 1, 128))))
    newcmp = ListedColormap(newcolors, name='OrangeBlue')
    cmap= newcmp
    
    cmap.set_bad('g', 1)
       
    im = plt.pcolormesh(data_to_plot[:,:], cmap = cmap, vmax =2, vmin =-2, edgecolor = 'black',linestyle ='--',lw =.01)
    plt.colorbar(im, aspect=40)
    plt.xlabel("Physical Properties",fontsize = 18)
    plt.ylabel("Residue Number",fontsize = 18)
    #setting up x axis labels
    num_categories = len(Properties)
    x_positions = range(num_categories)
    offset = 0.35
    plt.xticks([x_pos + offset for x_pos in x_positions],Properties, rotation = 90, fontsize =16)
    plt.yticks(ytix,fontsize =16)
    plt.title("FG Ins NNKx3 Physical Properties",fontsize = 19)
    plt.ylim(0,3)
    plt.xlim(0,10)
    
    fig.tight_layout()
    
    plt.savefig(name, dpi=1000, format='pdf')
    
    return


# %% [code]
fgpos_plot_physical_properties(fgpos_pp_preferences,'FGPeptideIns_PhysicalProperties.PDF')
```
